# Optogenetic dissection of transcriptional repression in a multicellular organism

**DOI:** 10.1101/2022.11.20.517211

**Authors:** Jiaxi Zhao, Nicholas C. Lammers, Simon Alamos, Yang Joon Kim, Gabriella Martini, Hernan G. Garcia

## Abstract

Transcriptional control is fundamental to cellular function. However, despite knowing that transcription factors can repress or activate specific genes, how these functions are implemented at the molecular level has remained elusive. Here we combine optogenetics, single-cell live-imaging, and mathematical modeling to study how a zinc-finger repressor, Knirps, induces switch-like transitions into long-lived quiescent states. Using optogenetics, we demonstrate that repression is rapidly reversible (*∼*1 minute) and memoryless. Furthermore, we show that the repressor acts by decreasing the frequency of transcriptional bursts in a manner consistent with an equilibrium binding model. Our results provide a quantitative framework for dissecting the *in vivo* biochemistry of eukaryotic transcriptional regulation.

## 1 Introduction

Throughout biology, transcription factors dictate gene expression and, ultimately, drive cell-fate decisions that play fundamental roles in development (*1*), immune responses (*2*), and disease (*3*). Achieving a quantitative and predictive understanding of how this process unfolds over time and space holds the potential both to shed light on the molecular mechanisms that drive cellular decision-making and to lay the foundation for a broad array of bioengineering applications, including the synthetic manipulation of developmental processes (*4–8*) and the development of therapeutics (*9*).

In recent years, great progress has been made in uncovering the molecular mechanism of transcription factor action through cell culture-based methods thanks to the emergence of a wide array of imaging techniques that can query the inner workings of cells in real time, often at the single molecule level (see, for example, (*10–18*)). However, progress has been slower in multicellular organisms, where a lack of comparable tools has limited our ability to query the dynamics of transcription factor function in their endogenous context. Indeed, while fixation-based methods, such as immunofluorescence staining, mRNA FISH, and various sequencing-based techniques represent powerful tools for investigating cellular decision-making in animals (*19–28*), these methods are mostly silent regarding the single-cell and single-gene dynamics of transcriptional control.

To move beyond these limitations, new experimental techniques are needed that provide the ability to quantify and manipulate input transcription factor concentrations *over time* in multicellular organisms while simultaneously measuring output transcriptional activity. Recently, we and others have developed new technologies to realize this goal through new molecular probes that allow for the direct measurement of protein (*29*), and transcriptional dynamics (*30, 31*) in single cells of living multicellular organisms, as well as optogenetic techniques for the light-based modulation of nuclear protein concentration *in vivo* (*32, 33*).

Here, we combine these technologies to study causal connections between the molecular players that underpin transcriptional control, shedding new light on the molecular basis of transcriptional repression in a developing animal. We use this platform to answer two key questions regarding the kinetic properties of repression. First, despite several studies dissecting repressor action at the bulk level (*23, 34–37*), it is not clear whether this repression is implemented in a graded or switch-like fashion at individual gene loci over time (Figure 1A, left). Second, the adoption of cellular fates—often dictated by repressors—has been attributed to the irreversible establishment of transcriptional states (*2, 38, 39*). However, it is unclear whether the action of repressors is itself reversible—such that sustained repressor binding is required to maintain gene loci in inactive transcriptional states—or if, instead, repression is *irreversible*—such that even transient exposure to high repressor concentrations is sufficient to induce long-lived transcriptional inactivity as a result of, for example, the accumulation of chromatin modifications. To answer these questions, in this work, we examine how the zinc-finger repressor Knirps drives the formation of stripes 4 and 6 of the widely studied *even-skipped* (*eve*) pattern during the early development of the fruit fly *Drosophila melanogaster* (Figure 1B) (*40–42*).

**Figure 1:**
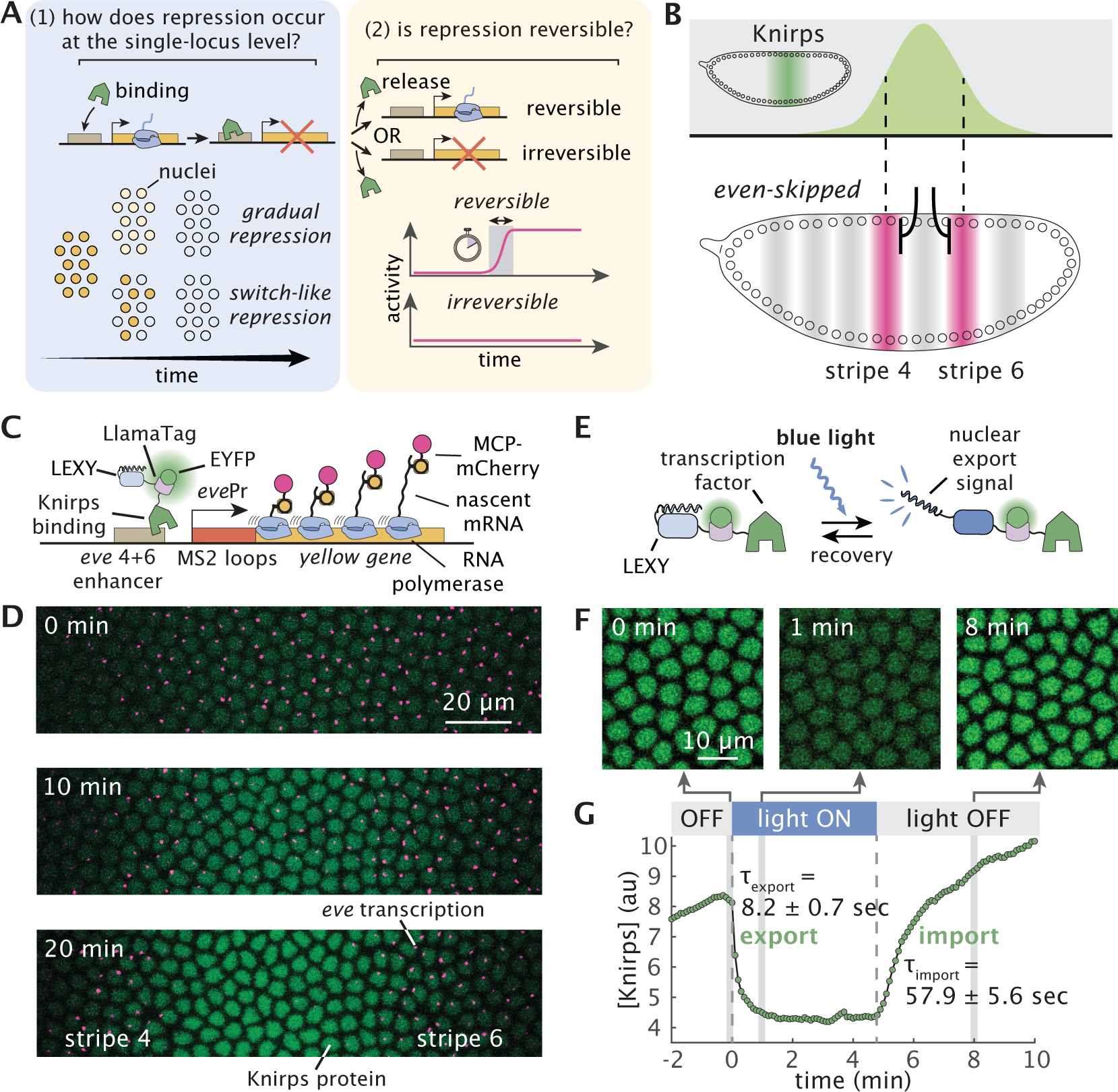
Combining optogenetics and live imaging enables dissection of single-cell repression dynamics in a developing animal. (**A**) Key questions regarding transcriptional repression. Left: Whether single-cell repression occurs in a gradual or switch-like fashion over time. Right: Whether repression reversible. (**B**) Knirps represses *even-skipped* (*eve*) stripes 4+6 transcription in the fruit fly embryo. Top: Knirps is expressed in a bell-shaped domain during early embryogenesis. Bottom: Knirps specifies the position and sharpness of the inner boundaries of *eve* stripes 4 and 6. (**C**) Two-color tagging permits the simultaneous visualization of input protein concentration and output transcriptional dynamics *in vivo*. Maternally deposited EYFP molecules bind to Knirps-LlamaTag, resulting in increased nuclear fluorescence, which provides a real-time readout of the nuclear protein concentration. Maternally deposited MS2 coat protein (MCP) binds to MS2 stem-loops in the nascent RNA formed by RNAP molecules elongating along the body of the *eve* 4+6 reporter construct leading to the accumulation of fluorescence at sites of nascent transcript formation. LEXY tag is also fused to Knirps to allow for optogenetic manipulation of its nuclear concentration. (**D**) Representative frames from live-imaging data. The embryo is oriented with the anterior (head) to the left. Green and magenta channels correspond to Knirps repressor and *eve* 4+6 transcription, respectively. When Knirps concentration is low, *eve* stripe 4+6 is expressed in a broad domain, which refines into two flanking stripes as Knirps concentration increases. (**E**) Optogenetic control of nuclear protein export. Upon exposure to blue light, the nuclear export signal within the LEXY domain is revealed. As a result, the fusion protein is exported from the nucleus. **(F)** Fluorescence images of embryos expressing the Knirps-LEXY fusion undergoing an export-recovery cycle. **(G)** Relative nuclear fluorescence of the repressor protein over time (*n* = 55 nuclei). Half-times for export and recovery processes are estimated by fitting the fluorescence traces with exponential functions.

## 2 Results

### 2.1 An optogenetics platform for dissecting single-cell repression dynamics in development

To measure Knirps protein concentration dynamics, we labeled the endogenous *knirps* locus with a LlamaTag, a fluorescent probe capable of reporting on protein concentration dynamics faster than the maturation time of more common fluorescent protein fusions (*29*). Further, we quantified the target transcriptional response using a reporter construct of the *eve* stripe 4+6 enhancer (*40*), where the nascent RNA molecules are fluorescently labeled using the MCP-MS2 system (*30, 31, 43*) (Figure 1C). The resulting nuclear fluorescence and transcriptional puncta provide a direct readout of input Knirps concentration and output *eve* 4+6 transcription, respectively, as a function of space and time (Figure 1D; Movie S1). Our data recapitulate classic results from fixed embryos (*44*) in dynamical detail: gene expression begins in a domain that spans stripes 4 through 6, subsequently refined by the appearance of the Knirps repressor in the interstripe region.

To enable the precise temporal control of Knirps concentration, we attached the optogenetic LEXY domain (*32*) to the endogenous *knirps* locus in addition to the LlamaTag (Figure 1C). Upon exposure to blue light, the LEXY domain undergoes a conformation change which results in the rapid export of Knirps protein from the nucleus (Figure 1E). Export-recovery experiments revealed that export dynamics are fast, with a half-time *<*10 seconds, while import dynamics are somewhat slower, with a half-time *∼*60 seconds upon removal of illumination (Figure 1F and G; Movie S2). These time scales are much faster than typical developmental time scales (*45*), allowing us to disentangle rapid effects due to direct regulatory interactions between Knirps and *eve* 4+6 from slower, indirect regulation that is mediated by other genes in the regulatory network. We established stable breeding lines of homozygous optogenetic Knirps flies, demonstrating that the protein tagged with both LEXY and LlamaTag is homozygous viable. Furthermore, our optogenetic Knirps drives comparable levels of *eve* 4+6 than wild-type Knirps (Figure S1). Thus, we conclude that our optogenetics-based approach represents an ideal platform for manipulating transcriptional systems to probe the molecular basis of gene regulatory control without significantly affecting the broader regulatory network and the developmental outcome this network encodes for.

### 2.2 Repressor concentration dictates transcriptional activity through all-or-none response

To understand how Knirps repressor regulates *eve* 4+6 expression, we first analyzed the temporal dynamics of Knirps-LlamaTag-LEXY (hereafter referred to simply as “Knirps”) concentration and *eve* 4+6 expression in the absence of optogenetic perturbations. We generated spatiotemporal maps of input repressor concentration and output transcription by spatially aligning individual embryos according to the peak of the Knirps expression domain along the anterior-posterior axis (Figure S2; Figure S3). These maps reveal a clear pattern: rising repressor concentrations coincide with a sharp decline in *eve* 4+6 activity at the center of the Knirps domain. Focusing on the central region of the Knirps domain (−2% to 2% of the embryo length with respect to the center of the domain), we observe a clear anti-correlation between Knirps concentration—which increases steadily with time—and the mean transcription rate, which drops precipitously between 10 and 20 minutes into nuclear cycle 14 (Figure 2A).

**Figure 2:**
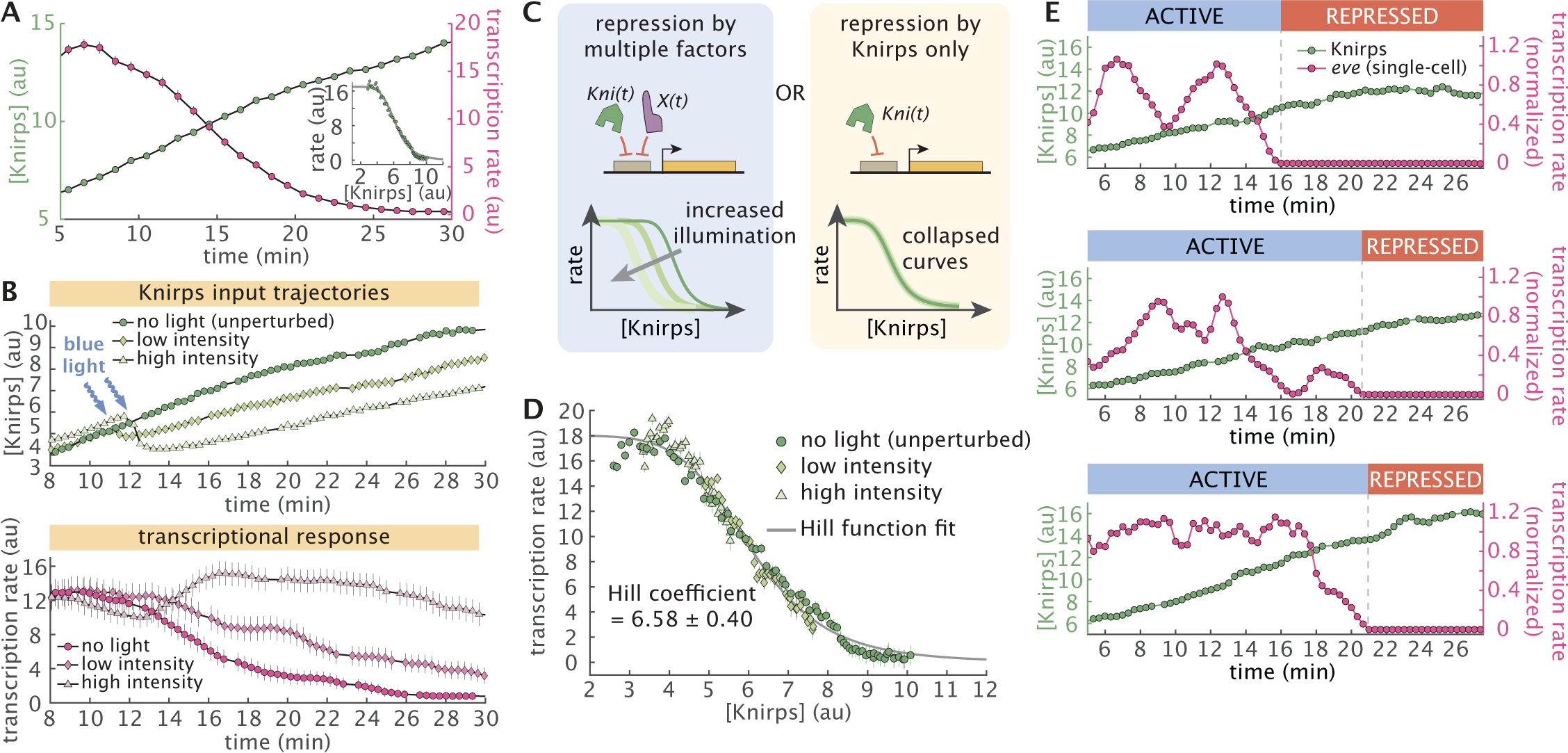
Knirps concentration dictates sharp, switch-like repression. (**A**) Average Knirps concentration (green) and *eve* 4+6 transcription (magenta) over time shows a clear anticorrelation. These dynamics are calculated by averaging the traces over a window of −2% to 2% along the anterior-posterior axis of the embryo and centered around the peak of the Knirps pattern (see Figure S2). Target transcription declines sharply as Knirps concentration increases. Inset panel shows the input-output relationship under this no light (unperturbed) condition. (**B**) Optogenetics allows for titration of protein concentration. Top panel shows the average Knirps concentration for three embryos, each under different illumination intensities. Bottom panel shows the corresponding trends in the *eve* 4+6 transcription rate. The illumination started around 12 minutes into nuclear cycle 14 and continued throughout the experiment. (**C**) To test whether Knirps is the only repressor whose concentration changes in the system, input-output functions under different illumination conditions can be compared. If there are multiple potentially unknown repressors at play (e.g. the *X* transcription factor in the figure), then each illumination level should lead to a different input-output function (left). However, if Knirps is the sole repressor, the input-output functions for each condition should collapse onto a single curve (right). (**D**) Average transcription rate as a function of Knirps concentration for each illumination condition (averaged over a window of −2% to 2% along the anterior-posterior axis). All three conditions follow the same trend, suggesting that Knirps is the only repressor regulating target transcription during this developmental stage. The input-output relationship is fitted with a Hill function resulting in a Hill coefficient of 6.36 (95% CI [6.08, 6.64]). (Averaged over *n* = 4 for no light, *n* = 4 for low intensity and *n* = 3 for high intensity embryos.) (**E**) Illustrative single-cell Knirps (green points) and transcriptional dynamics (magenta points) show that repression is switch-like at the single-cell level. Traces are normalized by their maximum transcription rate and smoothened using a moving average of 1 minute. (Error bars in A, B, and D indicate the bootstrap estimate of the standard error. *t* = 0 is defined as the onset of transcription in nuclear cycle 14. Transcription rate reflects the measured MS2 signal, which is an approximation of the *eve* mRNA production rate (*29, 30, 46*).)

We quantified the regulatory relationship implied by these trends by calculating the Knirps vs. *eve* 4+6 “input-output function”, which reports on the average transcription rate as a function of nuclear repressor concentration (inset panel in Figure 2A). This revealed a sharp decline in transcriptional activity across a narrow band of Knirps concentrations, suggesting that *eve* 4+6 loci are highly sensitive to nuclear repressor levels. This finding is consistent with previous observations that Knirps represses *eve* 4+6 (*47*), and with the discovery of multiple Knirps binding sites in the *eve* 4+6 enhancer region (Figure S4) (*48*). However, neither our endogenous measurements nor these previous studies can rule out the possibility that other repressors might also play a role in driving the progressive repression of *eve* 4+6 over the course of nuclear cycle 14. Indeed, by themselves, neither live imaging experiments (which are constrained to observing wild-type trends) nor classical mutation-based studies (which are subject to feedback encoded by the underlying gene regulatory network) can rule out the presence of other inputs.

Our optogenetics approach allows us to circumvent these limitations and search for regulatory inputs that impact *eve* 4+6 experiments, but are not directly observed in our experiments. Specifically, we used optogenetics to alter Knirps concentration dynamics over the course of nuclear cycle 14. Shortly after the beginning of the nuclear cycle, we exposed embryos to low and high blue light illumination, inducing moderate and strong reductions in nuclear Knirps concentration, respectively, which resulted in distinct transcriptional trends (Figure 2B; Figure S5; Movie S3). We reasoned that, because we are only altering Knirps concentration dynamics, the presence of other repressors dictating *eve* 4+6 activity together with Knirps should lead to distinct input-output curves across these different illumination conditions (Figure 2C, left). Conversely, if Knirps is the sole repressor driving the repression of *eve* 4+6 over time, the transcriptional input-output function should be invariant to perturbations of Knirps concentration dynamics (Figure 2C, right).

Comparing the *eve* 4+6 vs. Knirps input-output function for the unperturbed control (inset panel of Figure 2A) to that of optogenetically perturbed embryos (Figure 2D), we find that all three conditions collapse onto a single input-output curve, providing strong evidence that Knirps is the sole repressor of *eve* 4+6. Moreover, as noted above, we find that Knirps repression occurs in a sharp fashion: *eve* 4+6 loci transition from being mostly active to mostly repressed within a narrow band of Knirps concentrations. To quantify this sharp response, we fit a Hill function to the data in Figure 2D (gray line), which yielded a Hill coefficient of 6.58*±*0.40. Notably, this is comparable to Hill coefficients estimated for the Bicoid-dependent activation of *hunchback* (*20, 49, 50*); another canonical example of sharp gene regulation—in this case, of activation— during developmental patterning which relies on the presence of multiple binding sites for the transcription factor within the enhancer.

The input-output function in Figure 2D summarizes the average effect of repressor level on *eve* 4+6 expression, but it cannot alone shed light on *how* this effect is achieved in individual cells. Thus, we next investigated how this sharp *average* decrease in gene expression is realized at the single-cell level. We examined single-cell trajectories of Knirps repressor and corresponding *eve* 4+6 transcription. This revealed that the sharp population-level input-output function illustrated in Figure 2D is realized in an all-or-none fashion at the level of individual cells (Figure 2E; Figure S6). During this process, the gradual rise in Knirps concentration induces an abrupt, seemingly irreversible, transition from active transcription to a long-lived (or even permanent), transcriptionally quiescent state.

### 2.3 Rapid export of repressor reveals fast, reversible reactivation kinetics at the single-cell level

It has been shown that the activity of repressors can have different degrees of reversibility (*13, 51*). For example, recruitment of certain chromatin modifiers may silence the locus even if the initial transcription factor is no longer present (*13*). The single-cell traces in Figure 2E and Figure S6 *appear* to transition into an irreversible transcriptional quiescent state. However, since Knirps concentration keeps increasing after *eve* 4+6 expression shuts off, it is possible that repression is, in fact, reversible and that the observed irreversibility is due only to the monotonic increase of the repressor concentration over time.

To probe the reversibility of Knirps-based repression, we used optogenetics to induce rapid, step-like decreases in nuclear Knirps concentration (Figure 3A). Prior to the perturbation, the system was allowed to proceed along its wild-type trajectory until the majority of *eve* 4+6 loci at the center of the Knirps domain were fully repressed. Strikingly, when blue light was applied to export Knirps, we observed a widespread, rapid reactivation of repressed *eve* loci (Figure 3B and C; Movie S4). To probe the time scale of reactivation, we calculated the fraction of active nuclei as a function of time since Knirps export (Figure 3D, Figure S7). This revealed that *eve* loci begin to reactivate in as little as 1 minute following illumination. We obtain a reactivation time distribution from single-cell trajectories with a mean response time of 2.5 minutes (Figure 3E) and find that transcription fully recovers within 4 minutes of Knirps export (Figure 3D). Thus, Knirps repression is completely reversible.

**Figure 3:**
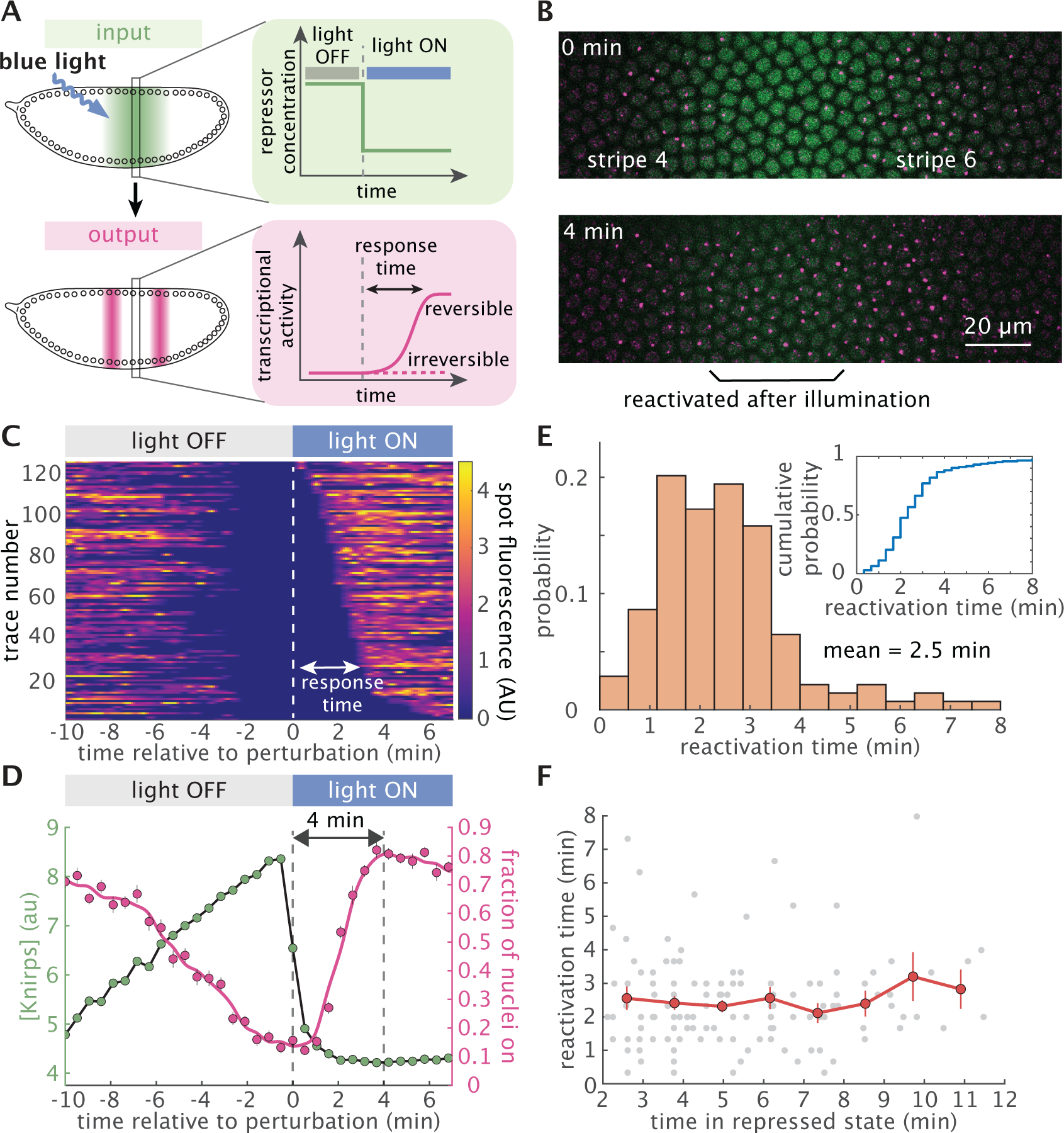
Knirps repression is rapidly reversible and memoryless. (**A**) Testing the reversibility of Knirps repression using a step-like optogenetic perturbation. Upon removal of Knirps repressor from the nucleus, transcriptional activity can remain repressed or recover, depending on whether repression is irreversible or reversible. (**B**) Snapshots from a movie before (top) and after (bottom) the optogenetic export of Knirps protein. Nuclei whose transcription was originally repressed by Knirps fully reactivate after 4 minutes of illumination. (**C**) Heatmap of single-cell reactivation trajectories sorted by response times. Response time is defined as the interval between the perturbation time and when the MS2 spots reappear. (**D**) Average repressor concentration (green) and the fraction of actively transcribing cells (magenta) before and after blue light illumination. We find that Knirps repression is rapidly reversible within 4 minutes. (*n* = 229 nuclei from 4 embryos, averaged over a −2% to 2% window along the anterior-posterior axis centered on the Knirps concentration peak). (**E**) Fast reactivation occurs with an average of 2.5 minutes. The reactivation response time is calculated as the interval between the perturbation and when a locus is first observed to resume transcription. (*n* = 139 nuclei from 4 embryos). Inset panel describes the cumulative distribution of reactivation times. To exclude gene loci that were transiently OFF due to transcriptional bursting or missed detections, we focused this analysis on gene loci that were silent for at least 2 minutes before perturbation. (**F**) Knirps repression is memoryless. Plot showing the reactivation response time of individual loci as a function of the time spent in the repressed state before optogenetic reactivation. The reactivation response time is independent of the repressed duration of the locus. (Error bars in D and F indicate the bootstrap estimate of the standard error.)

Previous studies have revealed regulatory “memory” wherein the repressive effect of certain repressors increases with longer exposure (*13*). Thus, we reasoned that prolonged exposure to high levels of a repressor could induce the accumulation of specific chemical or molecular modifications that prevent activator binding and, as a result, impede reactivation at the target locus, such as histone modifications (*52*). If this process is present, we should expect gene loci that have been repressed for a longer period before optogenetically triggering repressor export to require more time to reactivate. To test this hypothesis, we used the measured single-cell reactivation trajectories (Figure 3C) to calculate the average reactivation time as a function of how long cells had been repressed prior to Knirps export. Interestingly, our analysis reveals that the reactivation time has no dependence on the repressed duration (Figure 3F). This, combined with the fact that nearly all (97%) repressed gene loci reactivate upon Knirps export (inset panel in Figure 3E), argues against the accumulation of any significant molecular memory amongst repressed gene loci within the *∼*10 minute time scale captured by our experiments. Instead, it points to a model where repressor action is quickly reversible and memoryless.

### 2.4 Knirps acts by inhibiting the initiation of transcription bursts

One of the simplest model that can capture the reversible, memoryless transitions between active and inactive transcriptional states observed in Figure 3 is a two-state model, in which the gene promoter switches stochastically between periods of transcriptional activity (“bursts”) and periods of inactivity (*42, 46, 50, 53–57*). Here, the gene promoter switches between active (ON) and inactive (OFF) states with rates *k*_on_ and *k*_off_, and initiates RNAP molecules at a rate *r* while in the ON state (Figure 4A). Consistent with this model, our single-cell transcriptional traces show clear signatures of transcriptional bursting (see, e.g., top two panels of Figure 2E; Figure S6), suggesting that this two-state framework provides a viable basis for examining how Knirps regulates transcriptional activity at *eve* 4+6 loci.

**Figure 4:**
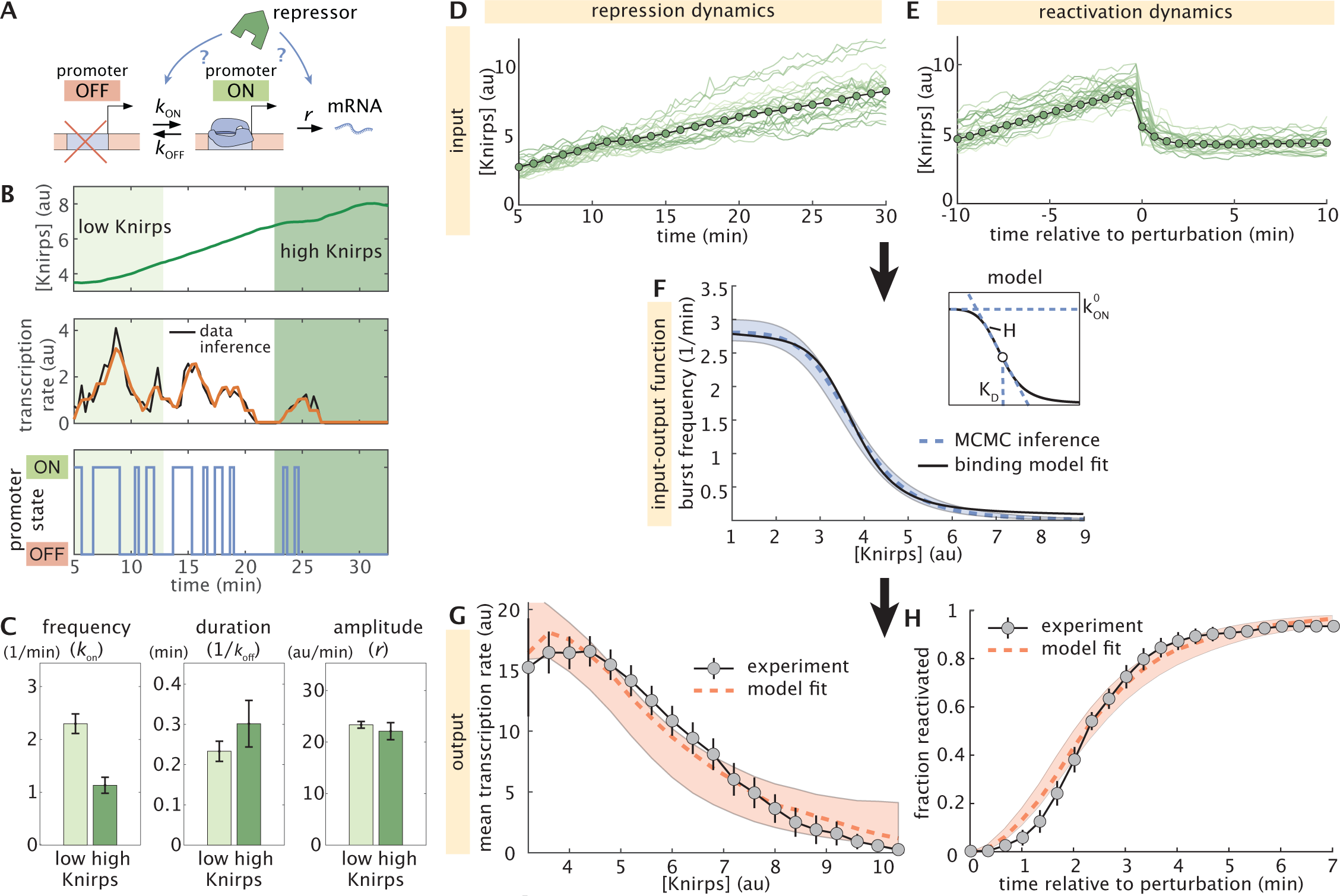
Knirps represses through instantaneous modulation of burst frequency. (**A**) Cartoon illustrating the two-state model of transcriptional bursting where a promoter can stochastically transition between active and inactive states. Knirps may regulate *eve* by altering any of the three kinetic rates in the model. (**B**) A representative experimental trace of Knirps protein (top) and transcription dynamics, along with the best fit (middle) and the corresponding sequence of inferred promoter activity states (bottom) returned by cpHMM inference. (**C**) Bar plots indicating cpHMM burst parameter inference results for *eve* 4+6 loci subjected to low ([Knirps]*≤* 4 au) and high ([Knirps]*≥* 6 au) Knirps concentrations. Inference reveals a two-fold decrease in the burst frequency, a moderate (30% though within error bars) increase in burst duration, and no notable change in the burst amplitude between low and high concentrations. (**D**-**H**) Summary of stochastic simulation methodology and results. First, we sample real single-cell Knirps concentration trajectories from (i) the three illumination conditions shown in Figure 2D and (ii) the reactivation experiments. (**D**) Illustrative individual (green lines) and average (green circles) nuclear Knirps concentration trajectories as a function of time in unperturbed embryos. **(E)** Individual and average nuclear Knirps concentrations before and after optogenetic export, which happens at time *t* = 0. **(F)** We take *k*_on_ to be a Hill function of Knirps concentration, with a shape that is determined by three microscopic parameters: 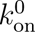, *K_D_*, and *H* (see inset panel and Equation 1). Given some set of microscopic parameters, we can plug Knirps concentration trajectories from (D) and (E) into the corresponding *k*_on_ input-output function to predict transcriptional outputs. The dashed blue curve indicates the input-output function for the burst frequency trend (*k*_on_) corresponding to the best-fitting set of microscopic parameters. Light blue shading indicates the standard error of the mean of the *k*_on_ input-output trend, as estimated by MCMC inference. To test the possibility that Knirps binding at the *eve* 4+6 enhancer, we fit a simple thermodynamic model to the trend revealed by our input-output simulations. The black line shows the best-fitting curve predicted by this molecular model. The binding model assumes 10 Knirps binding sites. We used the input-output function in (F) to generate a population of simulated MS2 traces that we used to predict. **(G)** the average fluorescence as a function of Knirps concentration and (**H**) the reactivation dynamics. Dashed red line indicates the prediction of the best-fitting model realization. Shaded red regions indicate standard deviation of the mean, as indicated by our MCMC inference. (Error bars in C reflect the standard error of the mean, as estimated from no fewer than 20 bootstrap burst inference replicates. The transcription rate is calculated from the measured MS2 signal, which is an approximation of the mRNA production rate (*29, 30, 46*).)

Within this model, the repressor can act by decreasing burst frequency (decreasing *k*_on_), by decreasing the duration of transcriptional bursts (increasing *k*_off_), by decreasing the burst amplitude (decreasing *r*), or any combination thereof as shown in Figure 4A. To shed light on the molecular strategy by which Knirps represses *eve* 4+6, we utilized a recently-developed computational method that utilizes compound-state Hidden Markov Models (cpHMM) to infer promoter state dynamics and burst parameter values (*k*_on_, *k*_off_, and *r*) from single-cell transcriptional traces as a function of Knirps concentration (Figure 4B) (*46*). We used data from all three illumination conditions (outlined in Figure 2B) and conducted burst parameter inference on 15-minute-long segments of MS2 traces.

To reveal burst parameter dependence on Knirps concentration, we grouped traces based on low ([Knirps]*≤* 4 au) and high ([Knirps]*≥* 6 au) Knirps concentrations (Figure 4B) and conducted cpHMM inference. We find that the repressor strongly impedes locus activation, decreasing the frequency of transcriptional bursts (*k*_on_) from 2.3 bursts per minute down to 1.1 burst per minute between low and high Knirps concentrations (Figure 4C, left panel). We also find a moderate (*∼* 30%) increase in the duration of transcriptional bursts between low and high Knirps concentrations; however this change is smaller than the uncertainty in our inference (Figure 4C, middle panel). Finally, we find no significant change in the burst amplitude as a function of Knirps concentration (Figure 4C, right panel). Thus, burst parameter inference indicates that Knirps represses *eve* 4+6 loci mainly by interfering with the initiation of transcriptional bursts. See Supplementary Text Section 1 and Figure S8 for additional cpHMM inference results.

To our knowledge, Figure 4C provides the first simultaneous measurement of transcription factor concentration and burst dynamics in a living multicellular organism. However, these results are, necessarily, a coarse-grained approximation of the true regulatory dynamics. This is because our cpHMM inference has an inherently low temporal resolution, reflecting averages taken across 15-minute periods of time and across large ranges of input Knirps concentrations. However, in principle, our live imaging data—which contains high-resolution time traces of both input repressor concentration dynamics and output transcriptions rates—should make it possible to move beyond these coarse-grained estimates to recover the true, *instantaneous* regulatory relationship between Knirps concentration and burst dynamics.

To answer these questions, we developed a novel computational method that utilizes stochastic simulations of single-cell transcriptional trajectories to test theoretical model predictions against our experimental measurements and uncover repressor-dependent burst parameter trends (Figure S9; Supplementary Text Section 2). Motivated by the cpHMM inference shown in Figure 4C, as well as finer-grained results shown in Figure S8, we allow both the burst frequency and the burst duration (but not the burst amplitude) to vary as a function of Knirps concentration. We assume a model in which these parameters are simple Hill functions of repressor concentration. For the burst frequency (*k*_on_), this leads to a function with the form

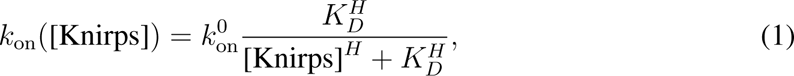

where 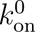 sets the maximum burst frequency value, the Hill coefficient *H* sets the sharpness of the response, and *K_D_* dictates the Knirps concentration midpoint for the transcriptional response, giving the repressor concentration where *k*_on_ drops to half its maximum value. Together, these “microscopic” parameters define an input-output function that directly links the burst frequency to Knirps concentration. As noted above, we also allow the burst duration to vary as a function of Knirps concentration (see Equation S2 and Supplementary Text Section 2.1 for further details). However we focus on *k*_on_ throughout the main text, since it is the only parameter that decreases as a function of Knirps concentration (and, thus, the only parameter that could drive *eve* 4+6 repression).

With our model defined, our procedure is as follows: we start by sampling real single-cell Knirps concentration trajectories from (i) the three illumination conditions shown in Figure 2D and (ii) the reactivation experiments shown in Figure 3 (Figure 4D and E, respectively). Then, we plug these Knirps trajectories into the input-output functions defined in Equation 1 (for burst frequency; see also Figure 4F) and Equation S2 (for burst duration). Next, given a set of microscopic parameters (e.g., *H*, *K_D_*, and *k*^0^ for Equation 1), we generate time-dependent burst parameter trends (Figure S9B). We then use these trends to simulate corresponding ensembles of MS2 traces (Figure S9C-F; see also Supplementary Text Section 2.1). We use these simulated MS2 traces to calculate, first, the predicted Knirps vs. *eve* 4+6 input-output function (Figure 4G) and, second, the predicted reactivation cumulative distribution function curve (Figure 4H). Finally, we compare these predictions to empirical measurements of the same quantities from our live imaging experiments (see Figure 2D and inset panel of Figure 3E). Through this process of simulation and comparison, each set of microscopic parameters used to calculate our predictions are assigned a fit score. We then use parameter sweeps and Markov Chain Monte Carlo (MCMC) (*58, 59*) to search for parameters that most successfully reproduced our live imaging results (see Figure S9E-G and Appendices 2.3 and 2.4).

As illustrated in Figure 4F, we find that the best-fitting model features a sharp *k*_on_ versus Knirps input-output function (*H* = 6.05 *±* 0.7). We also find that *k*_on_ has a relatively low *K_D_* of 3.7 au *±* 0.13 with respect to the range of Knirps concentrations experienced by *eve* 4+6 loci (see Figure 2B, bottom), which implies that gene loci have a low concentration threshold for Knirps repression. As a result of this low threshold, *eve* 4+6 loci are effectively clamped in the OFF state (*k*_on_ *≤* 0.1 bursts per minute) once the Knirps concentration exceeds 6 au, which happens about 12 minutes into nuclear cycle 14 for the average nucleus at the center of the Knirps domain (Figure 2B, bottom). See Figure S10 and Supplementary Text Section 2.5 for full inference results. Our findings also demonstrate that a simple two-state model in which Knirps represses *eve* 4+6 by decreasing the frequency of transcriptional bursts is sufficient to quantitatively recapitulate both the sharp decrease in the average transcription rate with increasing Knirps concentration (Figure 4G) and the kinetics of reactivation following Knirps export (Figure 4H).

Our simulation results also shed further light on the dynamics of *eve* reactivation following the step-like optogenetic export of Knirps protein from the nucleus (Figure 3A). From Figure 3E and F, we know that it takes approximately 2-4 minutes following Knirps export for MS2 spots to reappear in our live-imaging experiments. Yet this is the time scale for *detection*—for the amount of time it takes for genes to produce detectable levels of transcription and, hence, MS2 fluorescence—and thus likely overestimates the true *eve* 4+6 response time. So how fast is it really? Our model, which accounts for the fluorescence detection limit, predicts that *k*_on_ recovers to half of its steady-state value within 30 seconds of the start of the optogenetic perturbation (Figure S11). Furthermore, we predict that half of all gene loci switch back into the transcriptionally active (ON) state within 102 seconds (1.7 minutes). Thus, it takes fewer than two minutes for *eve* 4+6 loci to “escape” Knirps repression and re-engage in bursty transcription.

## 3 Discussion

Taken together, our results point to a model wherein the repressor acts upon the gene locus while it is transcriptionally inactive (OFF) to inhibit re-entry into the active (ON) state. Consistent with this picture, we find that the functional relation between *k*_on_ and Knirps concentration inferred by MCMC inference is well explained by a simple equilibrium binding model where the burst frequency is proportional to the number of repressor molecules bound at the 4+6 enhancer (solid black curve in Figure 4F; see Supplementary Text Section 3 for details).

Our *in vivo* dissection provides important clues toward unraveling the molecular basis of repressor action. We show that Knirps repression is switch-like (Figure 2), memoryless (Figure 3F), and rapidly reversible (Figure 3E). Another key point is that, although our model predicts that gene loci require 1-2 minutes to reactivate and enter the ON state following the optogenetic export of Knirps from the nucleus (Figure S11), the model assumes that the burst frequency itself responds *instantaneously* to changing Knirps concentration (see Equation 1, blue curve in Figure S11). While no reaction can truly be instantaneous, the success of this model in describing repression dynamics points to an underlying mechanism controlling the burst frequency that rapidly reads and responds to changing repressor concentrations, likely within a matter of seconds—a timescale that is consistent with the fast binding and unbinding dynamics reported for eukaryotic transcription factors (*60*). Lastly, the success of the two-state bursting model (Figure 4A) at recapitulating Knirps repression dynamics (Figure 4G and H) suggests that the same molecular process may be responsible for both the short-lived OFF periods between successive transcriptional bursts (see, e.g., the middle panel of Figure 4B) and the much longer-lived periods of quiescence observed in repressed nuclei (e.g., Figure 3C), and that there may be no need to invoke an “extra” repressor-induced molecular state outside of the bursting cycle (*61–63*).

Previous work has established that Knirps plays a role in recruiting histone deacetylase (*64*) and that Knirps repression coincides with increased histone density at target enhancers such as the one dissected here (*23*). This suggests a model in which the repressor modulates the longevity of the OFF state by tuning the accessibility of enhancer DNA, which would impact activator binding (*23*). It is notable, however, that the 1-2 minute reactivation time scales revealed (Figure 3; Figure S11) are faster than most chromatin-based mechanisms measured *in vivo* so far (*13, 51, 60, 65, 66*). This rapid reversibility, along with the memoryless nature of Knirps repression, indicates that whatever the underlying mechanism, Knirps binding at the locus is *necessary* in order to maintain the gene in a transcriptionally inactive state at the stage of development captured by our live imaging experiments. Interestingly, we found that the modulation of burst frequency by Knirps can be recapitulated by a simple thermodynamic model predicting Knirps DNA occupancy (black line in Figure 4F; see Supplementary Text Section 3 for further details). This suggests that the wide repertoire of theoretical and experimental approaches developed to test these models (see, for example, (*67*)) can be used to engage in a dialogue between theory and experiment aimed at dissecting the molecular mechanism underlying the control of transcriptional bursting.

Critically, none of these molecular insights would have been possible without the ability to measure and acutely manipulate input transcription factor concentrations in living cells. Thus, by building on previous works using the LEXY technology in different biological contexts (*32, 33, 68–70*), our work demonstrates the power of the LEXY system for simultaneously manipulating—and measuring—nuclear protein concentrations and the resulting output transcriptional activity. This capability can serve as a quantitative platform for dissecting gene-regulatory logic *in vivo*. Moreover, the LEXY system improves upon many previously reported methods of optogenetic control in embryos (*71–80*) (see Supplementary Text Section 4 for further discussions).

Looking ahead, we anticipate that our live imaging approach, along with the quantitative analysis framework presented in this work, will provide a useful foundation for similar *in vivo* biochemical dissections of how the transcription factor-mediated control of gene expression dictates transcriptional outcomes, opening the door to a number of exciting new questions relating to transcriptional regulation, cell-fate decisions, and embryonic development that span multiple scales of space and time.

## Acknowledgments

We would like to thank Jack Bateman, Augusto Berrocal, Gary Karpen, Kirstin Meyer, Brandon Schlomann, Max Staller, Robert Tjian, Meghan Turner, and Orion Weiner for their comments on the manuscript. We thank all the Garcia Lab members for inspiring discussions. NCL was supported by NIH Genomics and Computational Biology training grant (5T32HG000047-18), the Howard Hughes Medical Institute, and DARPA under award number N66001-20-2-4033. HGG was supported by the Burroughs Wellcome Fund Career Award at the Scientific Interface, the Sloan Research Foundation, the Human Frontiers Science Program, the Searle Scholars Program, the Shurl and Kay Curci Foundation, the Hellman Foundation, the NIH Director’s New Innovator Award (DP2 OD024541-01), NSF CAREER Award (1652236), an NIH R01 Award (R01GM139913) and the Koret-UC Berkeley-Tel Aviv University Initiative in Computational Biology and Bioinformatics. HGG is also a Chan Zuckerberg Biohub Investigator.

## Author contributions

Conceptualization: JZ, NCL, SA, YJK, HGG Methodology: JZ, NCL, SA, YJK, HGG Resources: JZ, NCL, SA, YJK, GM, HGG Investigation: JZ, NCL, HGG Visualization: JZ, NCL, HGG Funding acquisition: HGG Project administration: HGG Supervision: HGG Writing – original draft: JZ, NCL, HGG Writing – review & editing: JZ, NCL, SA, YJK, HGG

## Data and materials availability

All materials are available upon request. All data are available in the main text or supplementary materials. All code is available in this paper’s Github repository.

## Supplementary Materials

## Materials and Methods

Supplementary Text

Figures S1 to S12

Tables S1 to S4

Movies S1 to S4

References (*81–87*)

## Materials and Methods

### Cloning and Transgenesis

The fly lines used in this study were generated by inserting transgenic reporters into the fly genome or by CRISPR-Cas9 genome editing, as described below. See Table S1 for detailed information on the plasmid sequences used in this study.

### Creation of tagged *knirps* loci using CRISPR-Cas9

To tag endogenous the *knirps* locus with the EGFP-LlamaTag and LEXY modules, we used CRIPSR-mediated homology-directed repair with donor plasmids synthesized by Genscript. gRNA was designed using target finder tool from flyCRISPR (https://flycrispr.org), and cloned based on the protocol from (*81*). A yw;nos-Cas9(II-attP40) transgenic line was used as the genomic source for Cas9, and the embryos were injected and screened by BestGene Inc.

### Creation of *eve* 4+6 reporter

The *eve* 4+6 enhancer sequence is based on 800bp DNA segment described in (*47*). The *eve* 4+6 reporter was constructed by combining the enhancer sequence with an array of 24 MS2 stemloops fused to the *D. melanogaster yellow* gene (*29*). The eve4+6-MS2-Yellow construct was synthesized by Genscript and injected by BestGene Inc into *D. melanogaster* embryos with a ΦC31 insertion site in chromosome 2L (Bloomington stock #9723; landing site VK00002; cytological location 28E7).

### Transgenes expressing EYFP and MCP-mCherry

The fly line maternally expressing MCP-mCherry that is attached to a nuclear localization signal (chromosome 3) was constructed as described in (*29*). The fly line maternally expressing EYFP (chromosome 2) was constructed as previously described in (*82*). To simultaneously image protein dynamics using LlamaTags and transcription using MCP-MS2 system, we combined the vasa-EYFP transgene with MCP-mCherry to construct a new line (yw; vasa-EYFP; MCP-mCherry) that maternally expresses both proteins.

### Fly lines

To measure the Knirps pattern and corresponding *eve* 4+6 transcription simultaneously, we performed crosses to generate virgins carrying transgenes that drive maternal EYFP, MCP-mCherry, along with LlamaTag-LEXY tagged Knirps locus (yw; vasa-EYFP; MCP-mCherry/Knirps-LlamaTag-LEXY). These flies were then crossed with males having both the *eve* 4+6 reporter and LlamaTag-LEXY tagged Knirps locus (yw; eve4+6-MS2-Yellow; Knirps-LlamaTag-LEXY). This resulted in embryos homozygous or heterozygous for the tagged Knirps locus also carrying maternally deposited EYFP, MCP-mCherry, and a *eve* 4+6 reporter. Embryos homozygous for tagged Knirps can be differentiated from heterozygous embryos through a comparison of their nuclear fluorescence levels as shown in Figure S12. All the fly lines used in this work can be found in Table S2

### Embryo preparation and data collection

The embryos were prepared following procedures described in (*29, 30, 46*). Embryos were collected and mounted in halocarbon oil 27 between a semipermeable membrane (Lumox film, Starstedt, Germany) and a coverslip. Confocal imaging on a Zeiss LSM 780 microscope was performed using a Plan-Apochromat 40x/1.4NA oil immersion objective. EYFP and MCP-mCherry were excited with laser wavelengths of 514 nm (3.05 *µ*W laser power) and 594 nm (18.3 *µ*W laser power), respectively. Modulation of Knirps nuclear concentration was performed by utilizing an additional laser with a wavelength of 458nm, with laser power of 0.2 *µ*W (low intensity in Figure 2) or 12.2 *µ*W (high intensity in Figure 2 and Figure 3). Fluorescence was detected using the Zeiss QUASAR detection unit. Image resolution was 768 *×* 450 pixels, with a pixel size of 0.23 *µ*m. Sequential Z stacks separated by 0.5 *µ*m were acquired with a time interval of 20 seconds between each frame, except for the export-recovery experiment in Figure 1, in which we used 6.5 seconds.

### Image processing

Image analysis of live embryo movies was performed based on the protocol in (*30, 83*), which included nuclear segmentation, spot segmentation, and tracking. In addition, the nuclear protein fluorescence of the Knirps repressor was calculated based on the protocol in (*82*). The nuclear fluorescence of Knirps protein was calculated based on a nuclear mask generated from the MCP-mCherry channel. Knirps concentration for individual nuclei was extracted based on the integrated amount from maximum projection along the z-stack. The YFP background was calculated based on a control experiment and subsequently subtracted from the data.

### Predicting Knirps binding sites

To dissect Knirps binding to the *eve* 4+6 enhancer, we used Patser (*84*) with already existing point weight matrices (*85*) to predict Knirps binding sites. The predicted binding sites with scores higher than 3.5 are shown in Figure S4.

### Compound-state Hidden Markov Model

To obtain the inference results shown in Figure 4C, transcriptional traces were divided into 15 minute-long segments. Each trace segment was then assigned to an inference group based on the average nuclear Knirps concentration over the course of its 15-minute span. Trace segments with an average Knirps concentration of less than or equal to 4 arbitrary fluorescence units (au) were assigned to the “low” group and segments with a Knirps concentration greater than or equal to 6 au were assigned to the “high” group. Parameter estimates for each group were estimated by taking the average across 25 separate bootstrap samples of the “high” and “low” trace segment groups. Each bootstrap sample contained a minimum of 6,027 and 10,000 time points for the high and low groups, respectively. Inference uncertainty was estimated by taking the standard deviation across these bootstrap replicates. We used a model with two burst states (OFF and ON) and an elongation time of 140 seconds (equal to seven time steps; see (*46*)).

## Supplementary Text

## 1 Additional cpHMM inference results

In this section, we briefly describe additional cpHMM inference results. In addition to the binary inference results shown in Figure 4C that examine burst parameter values at high and low Knirps values, we also conducted finer-grained cpHMM inference runs, in which we queried burst parameter values across the full range of Knirps concentrations observed in our experiments. The plots in Figure S8 summarize our results. As with the results in the main text, this inference was conducted on 15-minute-long fragments of transcriptional traces. Multiple such fragments were generated from each transcription trace by sliding a 15-minute window along each and sampling in 1 minute increments. This produced a dataset of transcriptional “reads” that were then grouped by average Knirps concentration. In addition, we grouped transcriptional reads by experiment type (as defined in Figure 2B and D): no light (circles in Figure S8), low intensity (diamonds), and high intensity (squares).

We find that the inference results are consistent with the trends indicated in Figure 4C. We once again see that the burst frequency decreases with increasing Knirps concentration, though it is notable that the increased dynamic range of our inference reveals a more dramatic dependency, with burst frequency (*k*_on_) dropping by a factor of 6 across the range of concentrations examined (Figure S8A). Additionally, we see that the burst duration (1*/k*_off_) increases with increasing Knirps and that burst amplitude (*r*) remains roughly constant (Figure S8B and C). We note that, on its own, the Knirps-dependent increase in burst duration would actually lead to *activation*. Thus, although the burst duration exhibits Knirps-dependence, the burst frequency is the only parameter that is modulated in a manner consistent with the reduction in transcription as a result of repressor action.

However, while these findings paint a more detailed picture of how Knirps regulates transcriptional dynamics than the binary results presented in the main text, their resolution is nonetheless still limited by the fact that we must use 15-minute fragments for cpHMM inference. As a result, this approach is not suitable for recovering the true, instantaneous input-output functions that dictate how Knirps dictates burst parameter values. To make progress toward this goal, we developed a simulation-based computational framework for input-output function inference. We provide further details on this approach in the following sections.

## 2 Stochastic input-output simulations

Here we provide further details regarding the implementation of the simulation-based computational method that was utilized to produce the results featured in Figure 4F-H of the main text. Our aims in developing this method were two-fold: first, we sought to use our live imaging data to uncover burst parameter input-output functions and, second, we sought to assess whether a simple two-state model of transcriptional control based on our inference results in Figure 4C is *sufficient* to explain both the sharp input-output function (Figure 2D) and rapid reactivation dynamics (Figure 3D-E) revealed by our experiments.

### 2.1 Model specification

Our coarse-grained cpHMM burst inference results indicate that both burst frequency (*k*_on_) and burst duration (1*/k*_off_) vary as functions of Knirps concentration (Figure 4C). Accordingly, we employed a modeling framework in which both of these parameters vary as a function of Knirps concentration. Specifically, we model *k*_on_ and *k*_off_ as simple Hill functions of nuclear Knirps concentration (see inset panel of Figure 4F), such that:

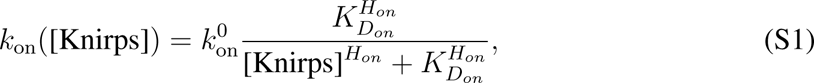

and

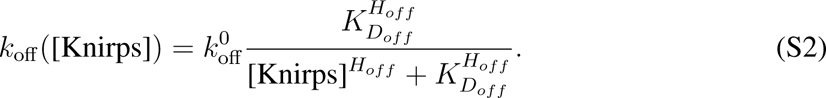

where 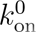 and 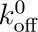 set the upper limits for on and off rates, respectively; where the Hill coefficient *H_on_* and *H_off_* set the sharpness of each parameter’s response to increasing Knirps concentration; and where *K_Don_* and *K_Doff_* dictate the half-max points for the *k*_on_ and *k*_off_ input-output curves. Finally, we assume that the burst amplitude, *r*, takes on a fixed value that does not vary as a function of Knirps concentration.

### 2.2 Stochastic simulations

We can use Equations S1 and S2 to generate simulated fluorescent traces with burst dynamics that vary as a function of nuclear Knirps concentration. To do this, we first sample real single-cell Knirps concentrations from (i) the three illumination conditions shown in Figure 2B and (ii) the reactivation experiments shown in Figure 3B-D (see also Figure 4D and E), and use these to generate time-dependent burst parameter trends. Figure S9A shows an illustrative time trace of Knirps concentration and panel Figure S9B shows the corresponding *k*_on_ (blue curve) and *k*_off_ (red curve) trends generated by plugging that trace into Equations S1 and S2. Note that the burst duration can be obtained simply by taking the inverse of the *k*_off_ trend. These burst parameter trends are used to simulate an ON/OFF promoter trajectory (Figure S9C), which, in turn, is used to generate a predicted MS2 trace (Figure S9D) with Knirps-dependent burst dynamics.

To simulate promoter trajectories with concentration-dependent burst parameters, we used a discrete implementation of the widely used Gillespie Algorithm (*86*), in which the promoter state is sampled with a time resolution of 1 second. We provide a brief overview of the approach here, and direct readers to the Github repository accompanying this work for further details regarding the algorithm’s implementation. Consider the time-varying burst parameter trends shown in Figure S9B, along with the simulated ON/OFF promoter trajectory in Figure S9C. At 11 minutes, we see that the promoter switches into the OFF state. In a standard Gillespie simulation with constant burst parameters, we would obtain the time until the next transition, *τ*_OFF_, by drawing a random sample from an exponential distribution with rate parameter *λ* = *k*_on_, such that

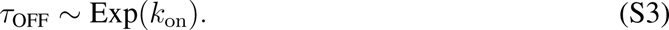

At time 11 + *τ*_OFF_, the promoter would then transition out of the OFF state and into the ON state.

Our case is more complicated, however, since *k*_on_ may change over time as the nuclear Knirps concentration changes. One simple way to capture this time-dependence is to adopt a discrete approach to promoter state simulations. In this approach, we designate some finite simulation time resolution, Δ*t*. Starting again at *t* = 11 minutes (with the promoter in the OFF state), the algorithm proceeds as follows:

1. Use Equation S1 to calculate *k*_on_ based off of the current Knirps concentration

2. Sample an expected jump time *τ*

if promoter is OFF, sample *τ* from an exponential distribution with rate parameter *k*_on_

else, sample *τ* from an exponential distribution with rate parameter *k*_off_

3. Compare *τ* to Δ*t*

if *τ ≥* Δ*t*: the promoter state remains unchanged

else, if *τ <* Δ*t*: change the promoter state (OFF to ON in our case)

4. Increment the time variable such that *t* = 11 + Δ*t*, and return to (1).

To understand why see this discrete rejection procedure for sampling the jump time *τ* is valid, consider the probability that the promoter remains in the OFF state for longer than *n* time steps (*P* (*τ*_OFF_ *> n*Δ*t*)). If we were sampling *τ*_OFF_ directly from the exponential distribution—as is the case for the standard Gillespie Algorithm—the probability of this outcome would be given by:

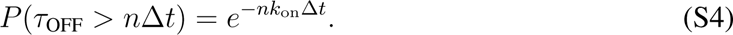

In our discrete rejection-based approach, the probability that *τ*_OFF_ *> n*Δ*t* is given by the joint probability that independently sampled values of *τ*, drawn at each iteration, are less than the sampling time resolution Δ*t*. The fact that each sample is independent means that the joint probability takes the form of a product:

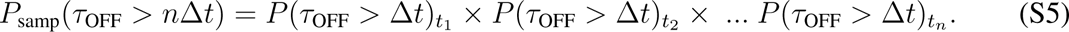

Simplifying, we see that the discretely sampled probability exactly equals the true probability

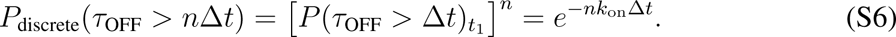

The main advantage of our discrete approach relative to the standard Gillespie Algorithm is that we are able to change the rate parameter (*k*_on_ or *k*_off_) at each sampling step to reflect changing Knirps concentrations. This leads to sampled jump time distributions of the form:

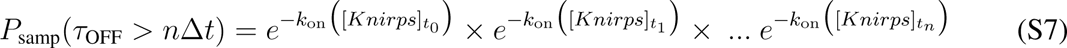

that reflect the effects of dynamic transcription factor concentrations.

Thus, by following our discrete sampling procedure, we obtain a discrete time trace of promoter activity, ***p***(*t*), that reflects time-dependent changes to the transition rates *k*_on_ and *k*_off_ due to changes in Knirps concentration. We set Δ*t* = 1 second, such that the resolution of our discrete sampling is significantly faster than the promoter burst dynamics being simulated (defined by *k*_on_ and *k*_off_; see Figure 4C). By enforcing this separation of timescales, we ensure that our discretely sampled time trace is a good approximation of a continuous Knirps-dependent trajectory.

Unlike *k*_on_ and *k*_off_, we assume that the initiation rates, *r*_0_ and *r*_1_, which encode the rate of Pol II initiation in the OFF and ON states, respectively, are Knirps-independent. Note that, for simplicity, we refer to *r*_1_ simply as “*r*” in the main text, and do not discuss results for *r*_0_ since *r*_0_ *≈* 0. Thus, to obtain a predicted time series of initiation rates, ***r*** from promoter states ***p***, we simply, set ***r*** = *r*_0_ for all time points when the promoter is OFF and ***r*** = *r*_1_ for all time points when the promoter is ON (see inset panel of Figure S9C). Finally, we obtain a predicted MS2 trace shown in Figure S9D by convolving ***r*** with the kernel *κ*_MS2_ Figure S9D, inset), which has the effect of taking a moving sum of past initiation rates over a time window defined by the time required for which nascent polymerase molecules remain on the gene body (set to 140 seconds throughout this work). This procedure also accounts for the finite amount of time needed for newly initiated Pol II molecules to transcribe the MS2 cassette and become fluorescent. We direct readers to Appendix D of (*46*) for further details.

### 2.3 Parameter sweeps

We used parameter sweeps to systematically test model performance across a broad range of plausible parameter values. As illustrated in Figure S9E, we performed a gridded sweep across 15 different values for *K_Don_* and *H_on_* from Equation S1. In addition we sampled 15 values each for *K_Doff_* and *H_off_* (not pictured) from Equation S2, making for a total of 15^4^ = 60625 distinct parameter combinations. The remaining parameters, namely 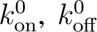, *r*_0_, and *r*_1_ were held fixed at their average values as calculated from the Knirps-dependent inference results shown in Figure S8A-C. Table S3 specifies the values and value ranges used for this procedure.

For each combination of parameter values, the procedure outlined in Figure S9A-D was used to generate ensembles of simulated fluorescent traces with realistic Knirps-dependent burst parameters using real experimental measurements of Knirps concentration over time (Figure S9F). We could then use these trace ensembles to calculate predictions for the fluorescence vs. [Knirps] input-output function and reactivation cumulative distribution function (CDF, Figure S9G). By comparing our model predictions to our experimental results (Figure S9G), it was possible to assess whether a given set of model parameters was sufficient to recapitulate these key features of Knirps repression.

We used the mean-squared error to assess model fits to the input-output function and reactivation CDF. In each case, deviations were normalized by the mean of the experimental curve to ensure comparable scaling between the fluorescence input-output errors (which are natively in arbitrary units) and CDF errors (which are probabilities). For the fluorescent input-output function (Figure 4G) this gives

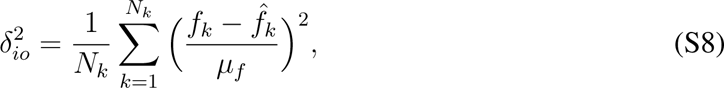

where *N_k_* is the number of Knirps concentration bins for which the average was calculated, *µ_f_* is the average fluorescence of the experimental curve in Figure 4G taken across all *N_k_* points, and where *f_k_* and *f*^^^*_k_* are the observed and predicted fluorescent values for Knirps concentration group *k*. Similarly, for the reactivation CDF we have

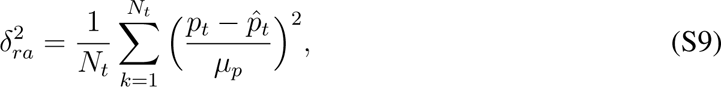

where *N_t_* is the number of time points post-reactivation that were considered, *µ_p_* is the average probability taken across the CDF in Figure 4H, and where *p_t_* and *p*^*_t_* are the observed and predicted fraction of reactivated nuclei at time point *t* post Knirps export.

We defined the total error in model fit as the weighted sum of 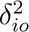 and 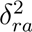, such that

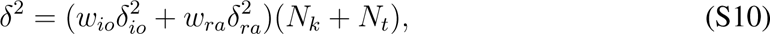

where the sum (*N_k_* +*N_t_*) up-weights *δ*^2^ according to the total number of data points considered, and where *w_io_* and *w_ra_* are weight parameters that tune the relative impact of 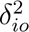 and 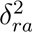 to the total loss, *δ*^2^. These weights can be adjusted to navigate tradeoffs between the minimization of input-output and reactivation CDF fitting loss. In our case, we find that values of *w_io_* = 1*/*4 and *w_ra_* = 3*/*4 lead to the best visual alignment between model predictions and experimental observations.

### 2.4 Estimating uncertainty bounds with MCMC

The parameter sweep procedure outlined above produced a *δ*^2^ estimate for each of the 60625 parameter combinations considered. In principle, the model realization corresponding to the lowest *δ*^2^ could be selected to obtain an approximate point estimate for the optimal *K_D_*, *H_on_*, *K_Doff_*, and *H_off_* values; however the parameter sweep results are not alone sufficient to obtain uncertainty bounds, nor do they provide insights into the remaining parameters not included in the sweep. To obtain this information, we employed Markov Chain Monte Carlo (MCMC) to sample the posterior distributions of our model parameters, conditional on our experimental data. MCMC is a widely used class of algorithms that are capable of efficiently sampling high-dimensional probability distributions (*58*).

As a first step in this process, we utilize information from the parameter sweeps to obtain parameter priors that are used to initialize and constrain MCMC sampling. To do this, we generate a weight vector, ***w***, comprised of terms with the form

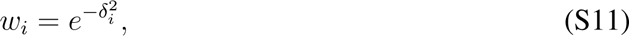

where 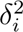 is the total loss from Equation S10 for the *i*th set of parameter values. If we assume that model errors are approximately Gaussian-distributed, then each *w_i_* can be interpreted as an unnormalized probability that is proportional to the likelihood of the data ***x*** (the input-output and reactivation curves) conditional on the *i*th parameter set ***θ_i_***:

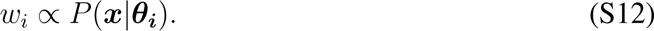

Moreover, from Bayes’ Theorem we have that

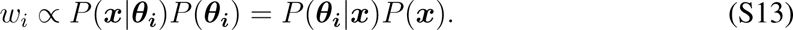

From here, we see that if we take a uniform prior across all ***θ_i_*** values (such that *P* (***θ_i_***) is a constant), then the weight *w_i_* will be proportional to the likelihood of each set of parameter values, conditional on the experimental data:

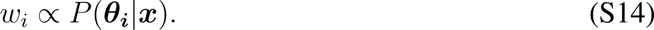

Motivated by this observation, we resampled the parameter values, ***θ***, surveyed in the parameter sweep according to the weight vector ***w***. This leads to a new set of parameter values, ***θ****^∗^*, where the frequency of a given parameter vector, ***θ_i_***, is proportional to its likelihood. As a result, the best-fitting parameter sets will appear more frequently in ***θ****^∗^*, and the worst-fitting are unlikely to appear at all. We calculate prior distributions for *K_Don_*, *H_on_*, *K_Doff_*, and *H_off_* (assumed to be Gaussian) by taking the mean and standard deviation of each parameters values across ***θ^∗^***. The prior distributions for 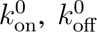, and *r*_1_ were initialized using the Knirps-dependent cpHMM inference results shown in Figure S8A-C. Specifically, the mean and standard deviation for 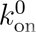 and 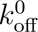 were estimated using the mean and standard deviations of the intercepts of the linear fits shown in Figure S8A and B, which we reasoned should provide reasonable estimates for the upper limit of each parameter. Given the lack of strong Knirps-dependence in the burst amplitude, the mean and standard deviation for the *r*_1_ prior were calculated by taking the mean and standard deviation of all cpHMM results shown in Figure S8C. The initiation rate when the system is in the OFF state, *r*_0_, was not subject to MCMC sampling, and was held fixed at its mean value from cpHMM inference. See Table S4 for the precise values used for each parameter prior.

With our prior distributions established, we conducted MCMC sampling to obtain estimates for the posterior distribution of each parameter. We conducted 24 independent MCMC simulations, each of which was run for 2500 total steps. We used standard Metropolis Hastings (*87*) updates during sampling. The procedure for each step was as follows:

1. At the *t*th step in the simulation, a new proposal for the parameter vector, ***θ^′^***, was generated by sampling from a multivariate normal distribution centered at the parameter values from the previous step, such that

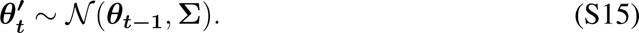 The covariance matrix, **Σ**, dictates how large or small the randomly proposed jumps tend to be relative to the previous parameter values. We assumed **Σ** to be a diagonal matrix and set each component, *σ_i_*, to be equal to 15% of the standard deviation of the corresponding parameter’s prior distribution.
2. Next, we used the proposed parameters, 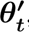, to simulate populations of MS2 traces and calculate predictions for the fluorescence vs. Knirps curve (Figure 4G) and reactivation CDF (Figure 4H) as outlined in the preceding sections.
3. We then calculated the total likelihood of the new parameters, defined as

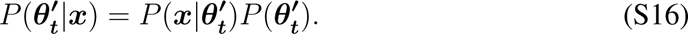 Here the first term on the right-hand-side is as defined in Equations S11 and S12, and functions to penalize proposals that produce curves that deviate too far from experimental measurements. The second component is the prior probability, and has the effect of penalizing proposals that deviate too far from our priors regarding parameter values.
4. Finally, we perform the standard Metropolis-Hastings move (*59, 87*). We calculate a probability, *p*, that takes the form

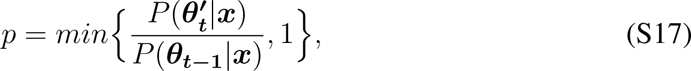

where *P* (***θ_t−_*_1_***|****x***) is the likelihood of the previous set of parameter values. Next we draw a random number, *z*, from the uniform distribution 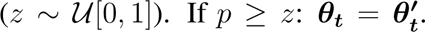. Otherwise: ***θ_t_*** = ***θ_t−_*_1_**.

### 2.5 Additional MCMC results

Figure S10 contains bivariate density plots and univariate histograms illustrating the results of MCMC sampling for each of the seven parameters examined. The results for the burst frequency (*k*_on_) are as quoted in the main text. We find that, like *k*_on_, *k*_off_ has a negative dependence on (*H_off_* = 3.2 *±*0.65). This translates to a burst duration that is predicted to *increase* as a function of increasing Knirps concentration (Figure S10C). On its own, this trend would *increase eve* 4+6 activity; however, this effect is dominated by the stronger Knirps-dependent decrease in *k*_on_, leading to a strong overall repressive effect (see Figure 4G). Additionally, our sampling returns a burst amplitude (*r*_1_) value of 21.6 *±* 1.9 au/min.

## 3 Implementation of the thermodynamic binding model

Here we provide a brief description of the theoretical underpinnings of the binding model that was used to generate the solid black curve in Figure 4F. The core assumption of this model is that *k*_on_ is proportional to the number of Knirps molecules bound to the locus, such that

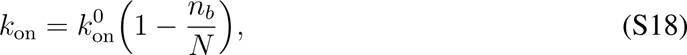

where 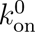 is the maximum burst frequency value (set to the 2.8 min*^−^*^1^ value returned by MCMC inference), *n_b_* is the number of Knirps molecules bound, and *N* is the total number of binding sites along the enhancer. Using PATSER scores for the *eve* 4+6 enhancer, we assess that there are 10 Knirps binding sites along the enhancer, such that *N* = 10 (see Figure S4). Thus, in this model *k*_on_ = 0 when *N* sites are bound and 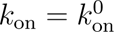 when 0 sites are bound.

Knirps-dependence enters into Equation S18 through *n_b_*, which should vary as a function of Knirps concentration. Note that, for ease of notation, we denote Knirps concentration by [*R*] (as opposed to [*Knirps*]) throughout this appendix. To model *n_b_*, we adopt the simple equilibrium chain model developed in (*60*). Briefly, this model assumes that all binding sites are identical, such that there are only *N* + 1 distinct binding states in which the enhancer can exist, ranging from 0 sites bound to all *N* sites bound. In this model, the Knirps concentration will induce a probability distribution over the set of possible binding states. Each state’s probability is given by

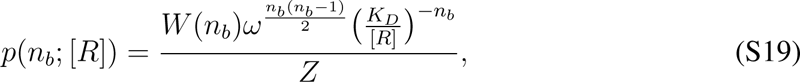

where *K_D_* is the dissociation constant for Knirps binding to specific sites at the locus. We assume that any pair of Knirps molecules can interact with a cooperativity factor *ω*. Given *n_b_* bound Knirps molecules, there are 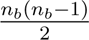 such pairwise interactions and, hence, a cooperativity contribution of 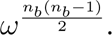. Further, *W* (*n_b_*) accounts for the number of different microscopic binding configurations that correspond to each macroscopic binding state (i.e., in how many different configurations can *n_b_* Knirps molecules be bound?):

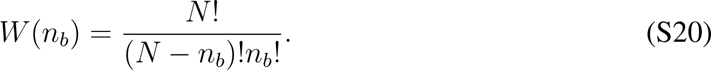

Lastly, we note that the denominator *Z* in Equation S19 is a normalizing factor equal to the sum of all *N* + 1 numerators. We direct the reader to Appendix C.3 of (*60*) for a detailed derivation of Equation S19.

We then use this expression for *p*(*n_b_*) to calculate the average expected number of bound Knirps molecules as a function of nuclear Knirps concentration, such that

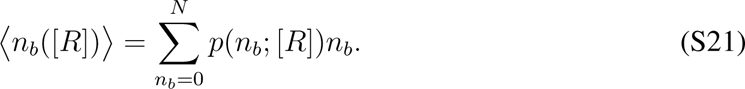

Finally, we plug Equation S21 into Equation S18 for the experimentally observed range of Knirps concentrations to produce a predicted burst frequency vs. Knirps input-output curve like the one shown in Figure 4F.

Using this approach, we conduct nonlinear least squares fitting to identify the values of *ω* and *K_D_* that best fit the blue *k*_on_ trend in Figure 4F that was returned by our MCMC inference. Our fit indicates that *K_D_* and *ω* values of 70.8 au and 1.9, respectively, produce the optimal fit. We find that the *k*_on_ trend generated by these parameter values (black in Figure 4F) is in close agreement with our MCMC inference result (blue curve), demonstrating that simple equilibrium binding could explain Knirps regulation of the burst frequency.

## 4 Comparison to other optogenetic approaches developed for multicellular organisms

In this work, we build on previous works using the LEXY technology (*32, 33, 68–70*) and demonstrate the power of the LEXY system for modulating protein dynamics inside developing embryos. The LEXY tag-based method addresses several key limitations faced by many previously reported methods.

First, some optogenetic tools are designed for specific signaling pathways (*71, 72, 76, 77, 79, 80*), and receptor (*74*) targets, and as a result, are not readily generalizable. In contrast, LEXY can be directly attached to any protein (though issues of genetic rescue (*69*) and its modulation strength (*33*) remain).

Second, many optogenetic tags do not act through concentration modulation, which makes it difficult to draw quantitative conclusions from the results. For example, the blue light-induced dimerization of *Arabidopsis* cryptochrome 2 (CRY2) controls downstream transcription by disrupting the function of the tagged protein through multimerization without affecting its concentration (*73, 75, 78*). On the other hand, LEXY controls transcriptional activity through direct modulation of the protein concentration within the nucleus, allowing for easy quantification and straightforward interpretation.

## Supplementary Figures

**Figure S1:**
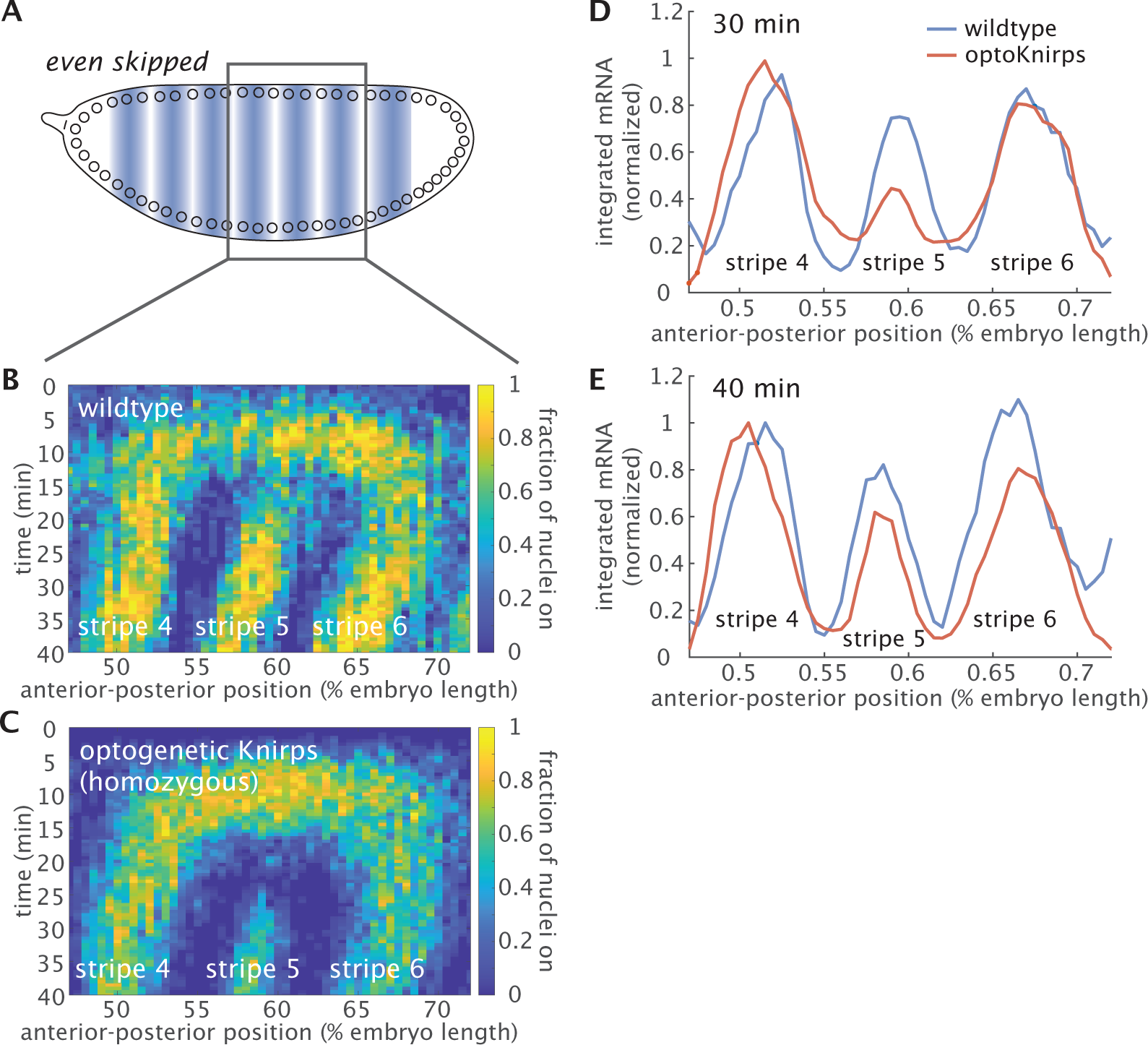
*even-skipped* expression under homozygous optogenetic Knirps (tagged with LEXY and LlamaTag) qualitatively recapitulates wild-type expression dynamics. (**A**) To understand whether and to what degree the *eve* expression pattern is impacted in the homozygous optogenetics Knirps background, we imaged the dynamics of a previously published eve-MS2-BAC reporter containing the full endogenous *eve* locus (*42*) in the wild-type and optogenetics Knirps backgrounds. (**B-C**) The expression pattern of *even-skipped* as reported by the fraction of detectable MS2 transcription spots is similar under wild-type Knirps (B) and optogenetics Knirps (C) except for a weaker stripe 5. (**D-E**) Comparison of the amount of mRNA present at 30 minutes into nuclear cycle 14 (as obtained by integrating the MS2 fluorescence signal) and at 40 minutes shows that stripe 5 expression is weaker under homozygous optogenetics Knirps at 30 minutes. The integration was performed assuming a mRNA half-time of 7 min. (D) Stripe 4 and 6 expression is slightly wider than under the wild-type condition at 30 min, suggesting that optogenetics Knirps is a slightly weaker repressor compared to the wild-type Knirps. (E) Stripe 5 expression continues to increase as it reaches a similar level compared to the wild-type around 40 minutes. The anterior-posterior position is aligned based on the center of stripe 5. The plots are normalized according to the peak of stripe 4 at 40 minutes and smoothened using a moving window of 1.5% range along the anterior-posterior axis. (Data from a single embryo is shown for each condition. *t* = 0 is defined as the onset of transc^3^ri^3^ption in nuclear cycle 14.)

**Figure S2:**
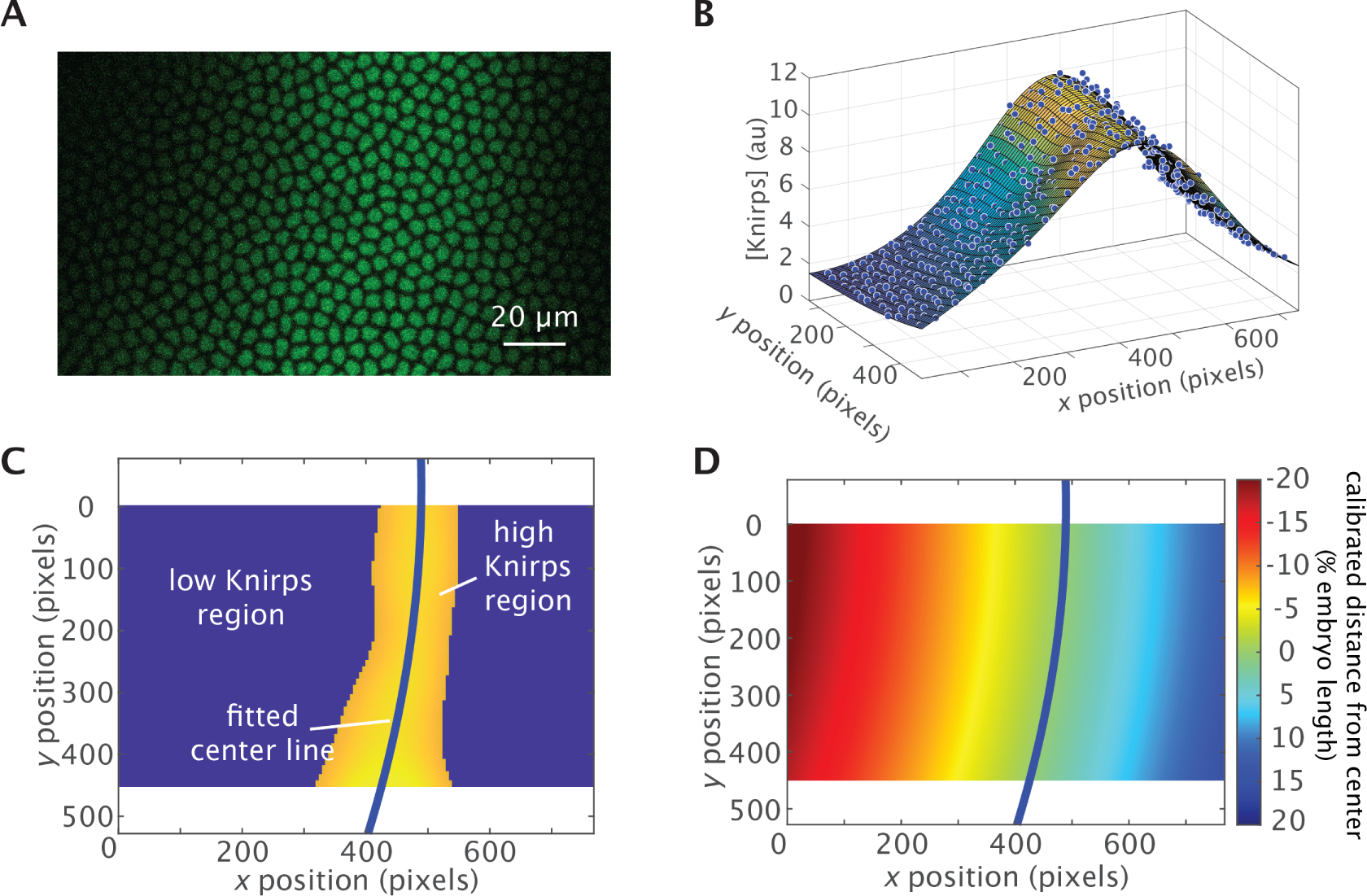
Nuclei position calibration based on Knirps expression pattern. The Knirps pattern of each individual embryo is used to align embryos along their anterior-posterior position axis. (**A**) Snapshot of the Knirps pattern used to calibrate nuclei position. (**B**) Extracted nuclear fluorescence is smoothed by local quadratic regression. (**C**) The region with high Knirps expression (yellow region) is extracted with a single threshold. Then, a quadratic function is fitted to the nuclei with high Knirps expression (yellow region) to extract the center line of Knirps expression (blue line). (**D**) Calibrated positions relative to the Knirps expression peak are calculated based on the distance to the extracted center line.

**Figure S3:**
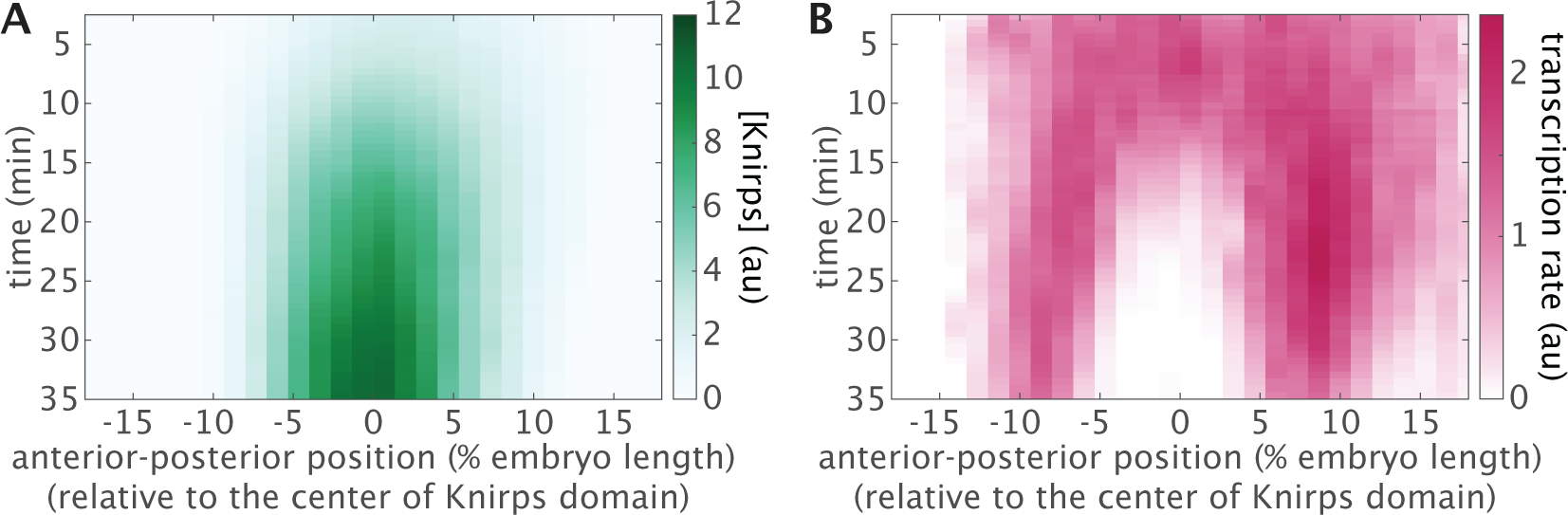
Spatiotemporal dynamics of Knirps protein and *eve* 4+6 transcription. Nuclei were binned based on their positions relative to the center of the Knirps domain (Figure S2, Materials and Methods) and their corresponding (A) Knirps protein concentration reported by LlamaTag fluorescence and (B) transcription reported by MS2 fluorescence were quantified over time.

**Figure S4:**
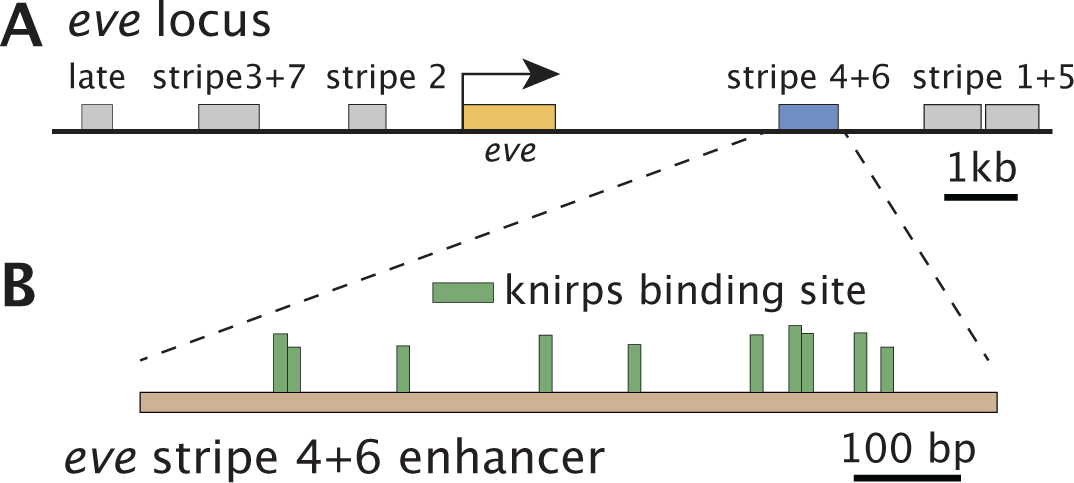
Predicted Knirps binding sites in the *eve* 4+6 enhancer. (A) The *eve* 4+6 enhancer is an 800 bp segment from the endogenous *eve* locus. (**B**) Ten Knirps binding sites are predicted within the *eve* 4+6 enhancer using PATSER (*84*) and Knirps position weight matrices from (*85*). Only binding motifs with PATSER scores higher than 3.5 are shown. The bar height of each binding site is proportional to the PATSER score.

**Figure S5:**
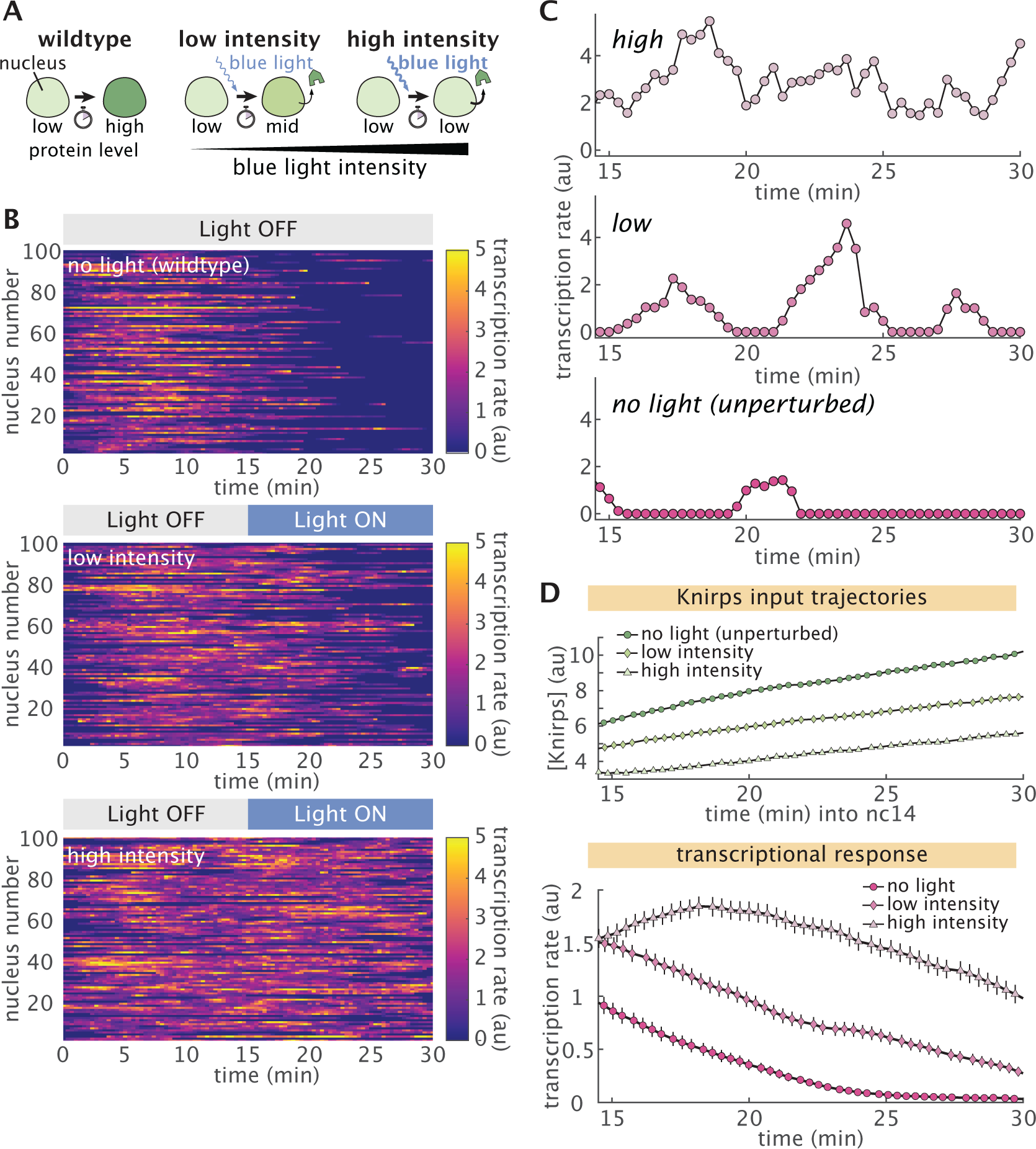
Repressor titration results in distinct transcriptional dynamics. (**A**) Optogenetic titration of protein concentration. Cartoon schematics for three different illumination conditions. Left: No illumination results in a negligible export of nuclear Knirps over time (green). Middle: Low dosage of blue light induces weak export of repressor from nuclei. Right: high intensity of blue light results in a strong export of repressor. (**B**). Single-cell traces for embryos with different Knirps export levels show distinct transcriptional dynamics. (**C**). Representative single-cell transcriptional dynamics under different illumination conditions show distinct responses. (**D**) Mean protein (top) and transcription rates (bottom) under different illumination conditions. Averaged over *n* = 4 (no light), *n* = 4 (low intensity) and *n* = 3 (high intensity) embryos. (Error bars in D indicate the bootstrap estimate of the standard error over multiple embryos.)

**Figure S6:**
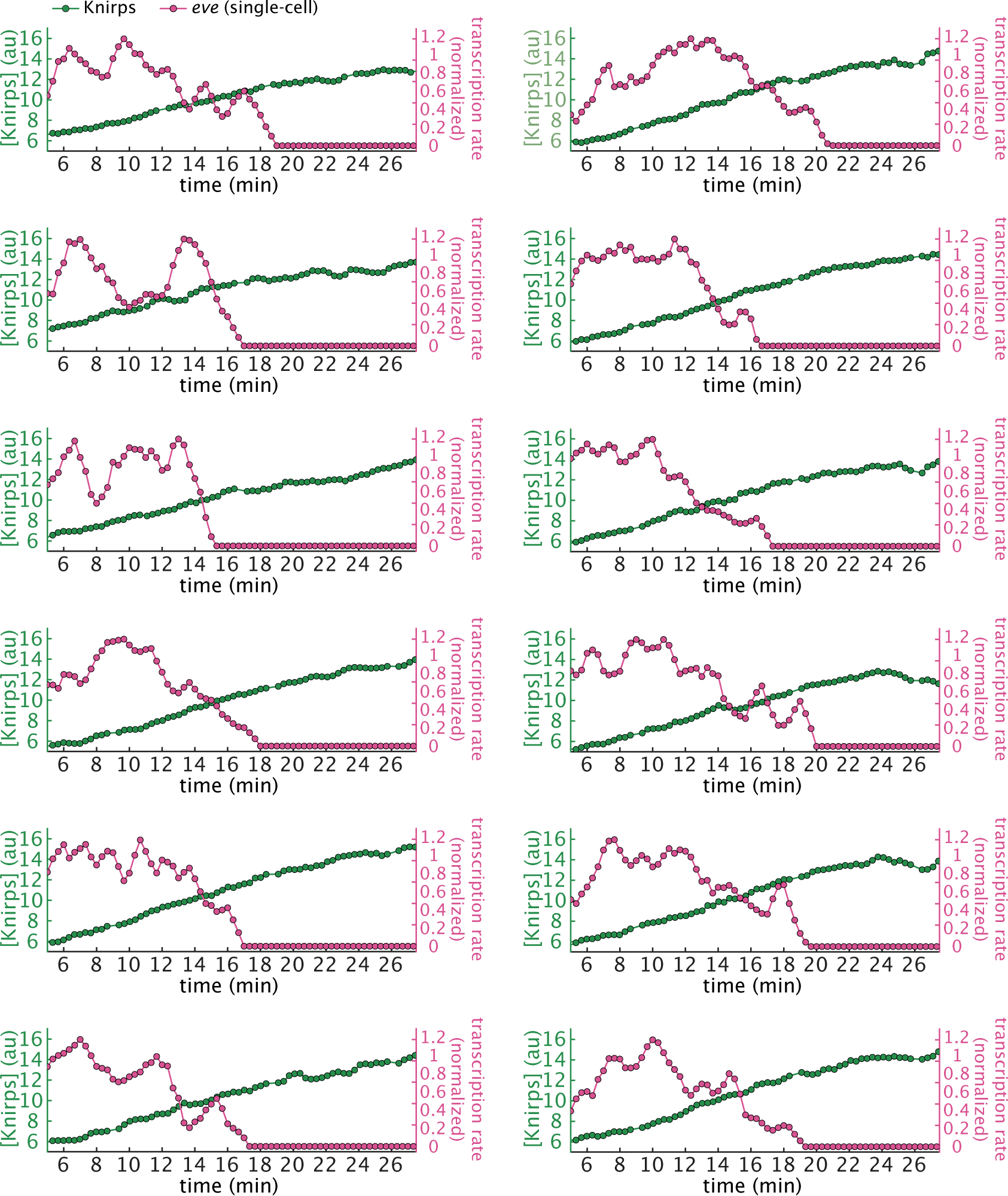
Example single-cell traces under no illumination. Single-cell input Knirps and output transcriptional dynamics traces show clear signs of transcriptional bursting, and that repression is switch-like. Traces are normalized by their maximum transcription rate and smoothened using a moving average of 1 minute.

**Figure S7:**
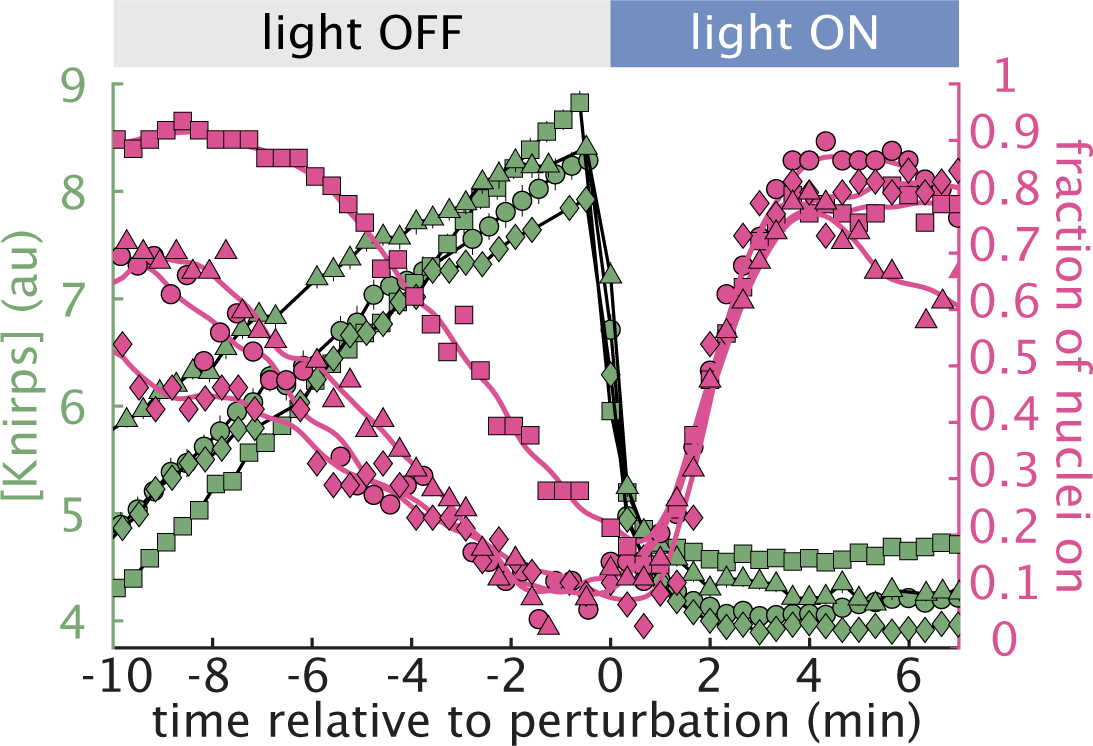
Response to Knirps perturbations is consistent across multiple embryos. Plot showing input Knirps concentration and output transcirptional activity for four individual embryos. All embryos display similar responses to Knirps export upon light exposure. Each marker shape corresponds to one embryo. (Error bars indicate the bootstrap estimate of the standard error.)

**Figure S8:**
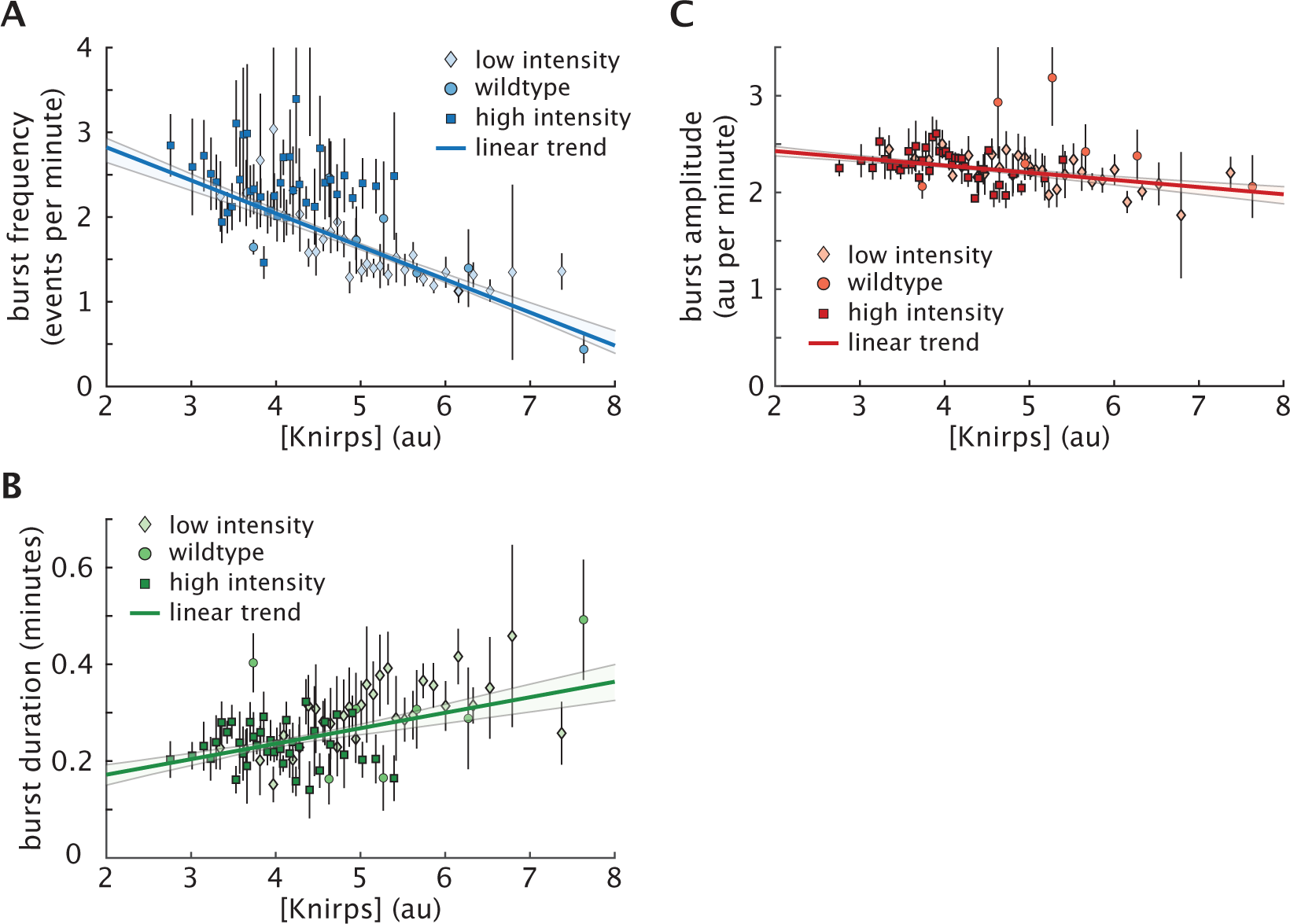
Full cpHMM inference results of Knirps-regulated transcriptional bursting. **(A)** We find that the burst frequency (*k*_on_) decreases significantly as a function of Knirps concentration. **(B)** We also find a moderate increase in burst duration (1*/k*_off_) with Knirps concentration, **(C)** while burst amplitude (*r*) remains approximately constant. Lines in A, B and C indicate the best linear fit to data. Circles, diamonds, and squares indicate data points from no light (unperturbed), low illumination, and high illumination experiments, respectively, as described in Figure 2B.

**Figure S9:**
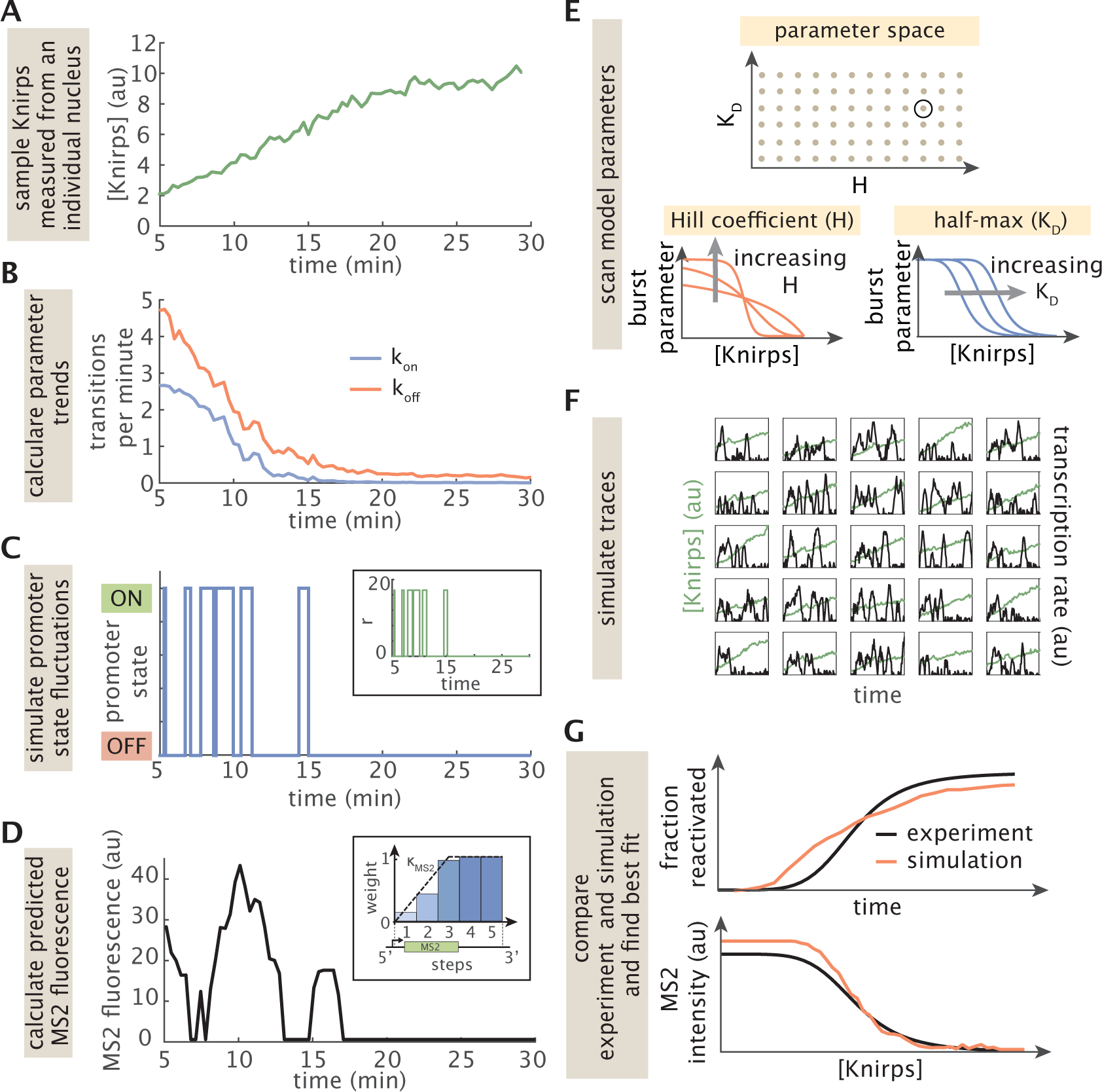
A computational framework for Knirps-dependent stochastic simulations. **(A-D)** Schematic showing process for simulating stochastic transcription time traces. **(A)** We first sample an empirical time trace of Knirps concentration from a nucleus in our live imaging dataset. **(B)** Next, we plug this Knirps trace into the input-output functions for *k*_on_ (Equation S1) and *k*_off_ (Equation S2) to generate time-dependent burst parameter trends. **(C)** We then use a discrete implementation of the Gillespie Algorithm to simulate a stochastic time-series of promoter activity that reflects the time-dependent parameter trends. Inset panel shows corresponding initiation rate time series. **(D)** Finally, we use this promoter time series to calculate the predicted MS2 fluorescence at each time point. We assume an initiation rate of 21.5 au when the promoter is in the ON state and a basal rate of 0.6 au when the promoter is OFF. **(E-G)** Schematic illustrating the parameter sweep algorithm. **(E)** We use a simple gridded search to sweep a broad space of values for key parameters in Equations S1 and S2. Cartoon illustrates case for a 2D search for *k*_on_-related parameters. In reality, we also scan the analogous *k*_off_ parameters, leading to a 4D gridded search. For each iteration of the sweep algorithm, we select a new combination of parameters (black circle in top panel). **(F)** Then, we use the process illustrated in A-D to simulate an ensemble of MS2 traces that reflect these parameter values. We generate one simulated MS2 trace for each experimental Knirps trace in our dataset **(G)** Finally, we use these simulated traces to calculate dynamics of the fraction of reactivated and MS2 fluorescence as a function of Knirps concentration for comparison with our experimental results. The mean squared error is used to assess agreement between prediction and experimental data and to identify the set of microscopic parameters that best describes the data.

**Figure S10:**
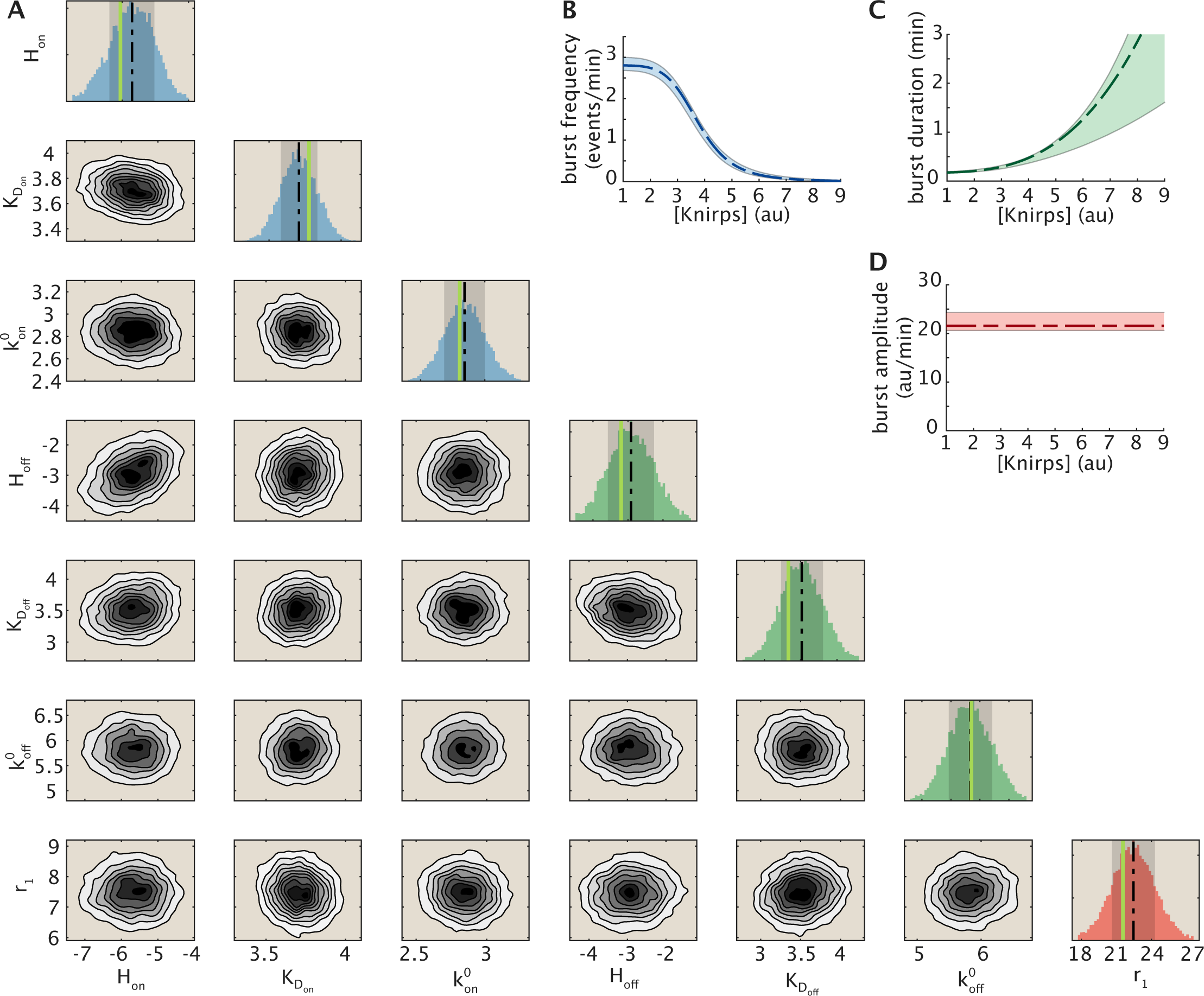
Full MCMC results for stochastic input-output model parameters. (**A**) Univariate and bivariate density plots. Vertical green lines in histograms indicate the mean parameter value taken across the 25 best-fitting model realizations. Dashed black lines indicate average parameter values taken across all MCMC samples; i.e. the full distribution shown in each histogram. Shaded regions in histograms indicate 1 standard deviation above and below the mean. (**B**) Inferred trends for the burst frequency (*k*_on_), **(C)** burst duration (1*/k*_off_) and **(D)** burst amplitude (*r*_1_). *k*_off_ was modeled as a Hill function of Knirps (see Equation S2) and *r* was assumed to be invariant relative to Knirps concentration.

**Figure S11:**
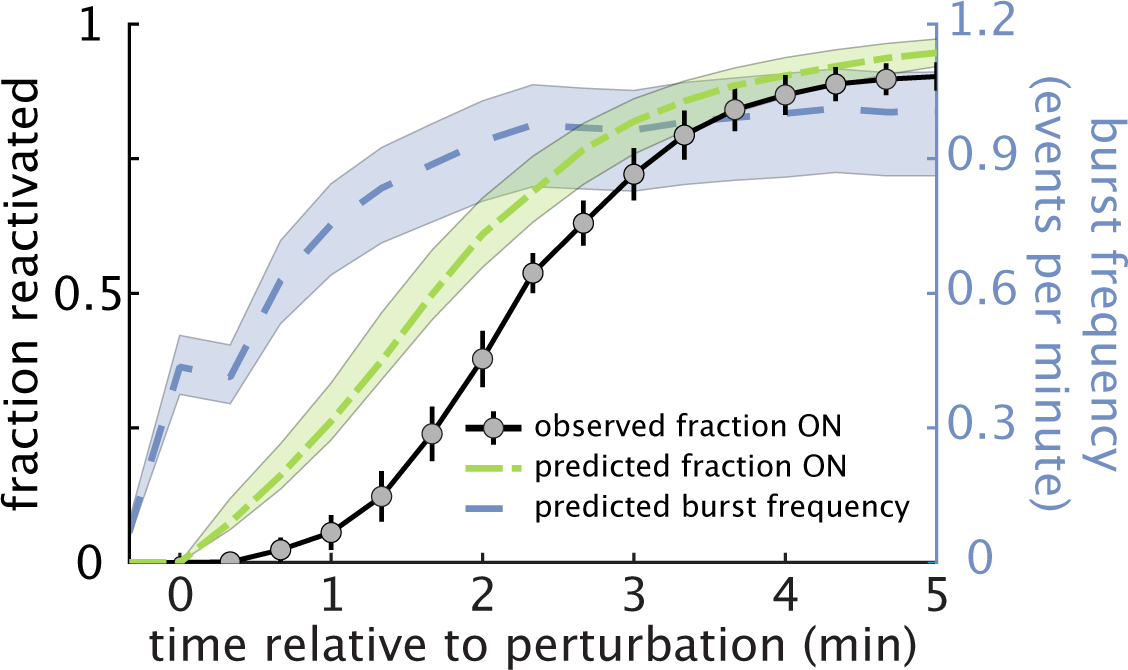
Model predictions for *eve*4+6 reactivation dynamics following Knirps export. Blue curve shows the predicted recovery of burst frequency (calculated from Equation S1) following the optogenetic perturbation of Knirps. The green curve indicates the corresponding cumulative fraction of loci that are predicted to have reentered the ON state as a function of time since the perturbation. Black curve is identical to the one shown in Figure 4H and corresponds to the measured fraction of loci that have reentered the ON state. We observe a lag between the cumulative fraction of ON loci and the experimentally observed fraction because recently reactivated gene loci typically require multiple time steps to accumulate sufficient fluorescent MS2 signal in order to be experimentally detected.

**Figure S12:**
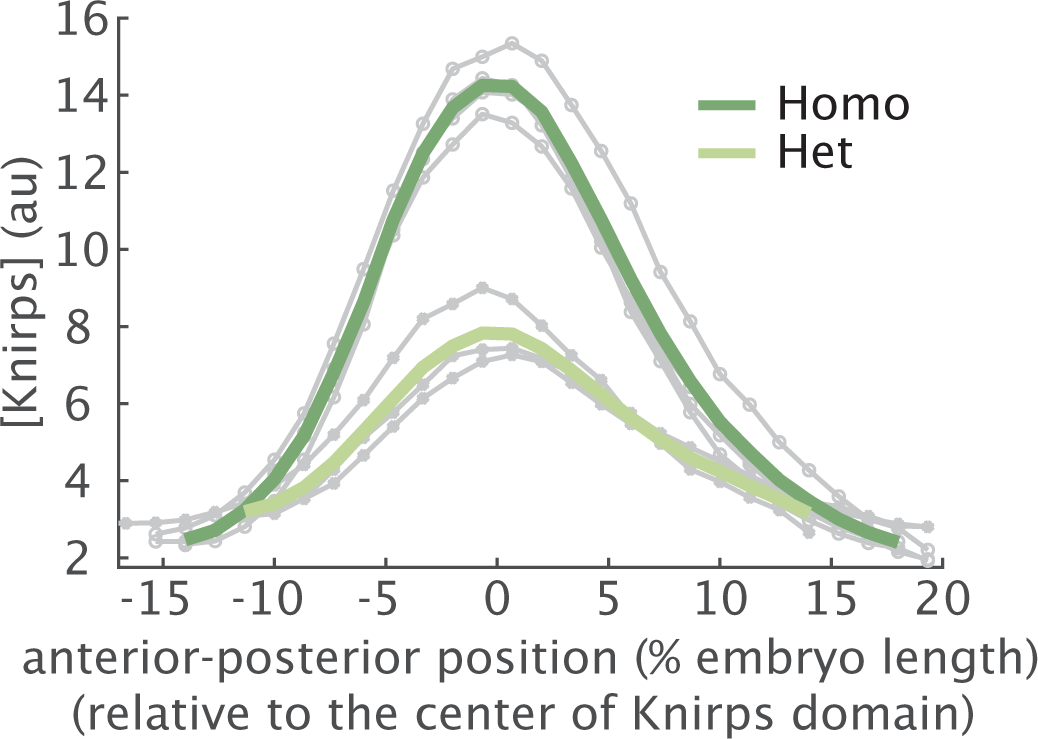
Distinguishing homozygous from heterozygous embryos. Homozygous embryos (*n* = 4) can be easily distinguished from heterozygous embryos (*n* = 3) by comparing Knirps concentration at 30 min into nc14.

## Supplementary Tables

**Table S1:** List of plasmids used in this study.

| **Name** | **Function** |
| --- | --- |
| pCasper-vasaPr-EYFP | P-element insertion plasmid for vasa promoter driven EYFP |
| pBPhi-eve4+6-evePr-MS2-Yellow | *eve* 4+6 reporter |
| pHD-Kni-LlamaTag-LEXY-dsRed | Donor plasmid for Knirps-LlamaTag-LEXY CRISPR knock-in fusion |
| pU6-3-gRNA-Knirps-1 | guide RNA 1 for Knirps-LlamaTag-LEXY CRISPR knock-in fusion |
| pU6-3-gRNA-Knirps-2 | guide RNA 2 for Knirps-LlamaTag-LEXY CRISPR knock-in fusion |

**Table S2:**
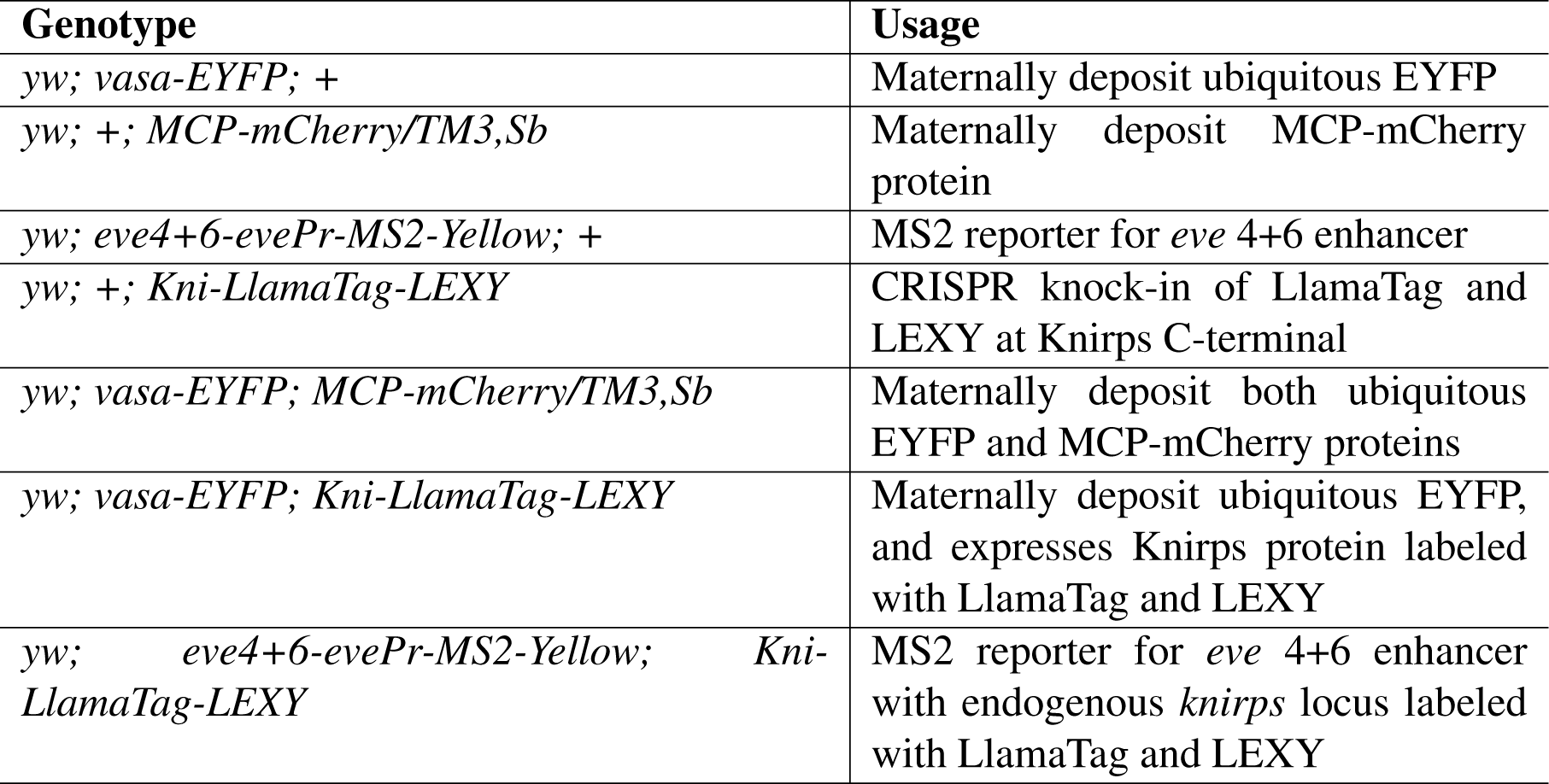
List of fly lines used in this study.

**Table S3:** List of parameter ranges used for parameter sweeps. Brackets denote inclusive ranges. Parameters with a single value appearing in the “range” column were held fixed during the sweeps. Parameters with two values were sampled at 15 equally spaced points bounded by the values indicated in the brackets.

| Parameter | Range |
| --- | --- |
| burst frequency Hill Coefficient ($H_{\text{ON}}$) | [3.15, 12.6] |
| burst frequency half-maximum ($K_{D\text{ON}}$) | [2.5, 10.2] (au) |
| max burst frequency ($k_{\text{on}}^0$) | 2.85 (events per min) |
| off rate Hill Coefficient ($H_{\text{OFF}}$) | [0, 4] |
| off rate half-maximum ($K_{D\text{ON}}$) | [2, 6] (au) |
| max off rate ($k_{\text{off}}^0$) | 5.81 (events per min) |
| ON state initiation rate ($r_1$) | 22.76 (au per min) |
| OFF state initiation rate ($r_0$) | 0.6 (au per min) |

**Table S4:** List of parameter priors used for MCMC sampling.

| Parameter | Prior distribution |
| --- | --- |
| burst frequency Hill Coefficient ($H_{\text{ON}}$) | $\mathcal{N}(5.7, 0.8)$ |
| burst frequency half-maximum ($K_{D\text{ON}}$) | $\mathcal{N}(3.7, 0.15)$ (au) |
| max burst frequency ($k_{\text{on}}^0$) | $\mathcal{N}(2.84, 0.17)$ (events per min) |
| off rate Hill Coefficient ($H_{\text{OFF}}$) | $\mathcal{N}(3.1, 0.8)$ |
| off rate half-maximum ($K_{D\text{ON}}$) | $\mathcal{N}(3.5, 0.3)$ (au) |
| max off rate ($k_{\text{off}}^0$) | $\mathcal{N}(5.8, 0.4)$ (events per min) |
| initiation rate ($r_1$) | $\mathcal{N}(22.8, 2.1)$ (au per min) |

## Supplementary Movies

**Movie S1 Full movie for repression without perturbation.** Knirps concentration is indicated in green. Active *eve* 4+6 loci appear in magenta. Timestamp indicates minutes since the start of nuclear cycle 14.

**Movie S2 Full movie demonstrating optogenetic manipulation of protein concentration.** Knirps concentration is indicated in green. Timestamp indicates time in minutes relative to the optogenetic perturbation.

**Movie S3 Full movie demonstrating optogenetic titration of protein concentration.** Panels correspond to the three illumination conditions illustrated in Figure 2B. Knirps concentration is indicated in green. Active *eve* 4+6 loci appear in magenta. Timestamp indicates minutes since the start of nuclear cycle 14.

**Movie S4 Full movie showing optogenetic export of repressor protein.** Knirps concentration is indicated in green. Active *eve* 4+6 loci appear in magenta. Timestamp indicates time in minutes relative to the perturbation.

